# Osteocyte Nicotinic Acetylcholine Receptors Impact Bone Mechanoadaptation in a Sexually Dimorphic Manner

**DOI:** 10.1101/2023.10.01.556129

**Authors:** Macy Mora-Antoinette, Andrea Garcia-Ortiz, Mariam Obaji, Alexander Saffari, Melia D. Matthews, Murtaza Wasi, Karl J. Lewis

## Abstract

Recent evidence suggests acetylcholine has a positive influence on bone mechanotransduction. Osteocytes express components for nicotinic acetylcholine receptors (nAChRs), which are known for mediating calcium signaling and may impact mechanosensitivity. Here, we use novel fluorescent imaging approaches to provide the first evidence of direct interaction between osteocytes and cholinergic nerve fibers in cortical bone *in vivo*. Moreover, we show that osteocytes are functional targets of cholinergic signaling for bone mechanoadaptation. We report sexually dimorphic patterns in bone structure and mechanobiology based on nAChR function. In females, osteocyte mechanosensitivity was decreased at small force magnitudes and tissue level deficits were recovered with anabolic loading. In males, osteocyte mechanosensitivity was increased in some groups and anabolic loading had very little effect on overall tissue architecture. This work establishes a new signaling paradigm wherein osteocytes interface with cholinergic nerves and bone mechanotransduction is regulated by osteocyte cholinergic signaling in a sexually dimorphic way.

## Introduction

Bone is an exquisitely mechanically sensitive tissue, capable of adapting its architecture and mass to maintain a structurally sound but energetically efficient form. Harold Frost proposed the idea that bone cells actively detect their strain environment to shape and maintain bone mass accordingly(*1*–*3*). He posited that if bone experienced prolonged periods of mechanical deformation below or above a set-point value, the mineralized matrix would be resorbed or deposited, respectively. Indeed, peak healthy *in vivo* deformations for a broad range of species are tightly conserved between 150-3000 με(*4*), and aberrant loading results in anabolic bone growth in the area of maximal bending(*5*). In some bone metabolic diseases, the relationship between tissue deformation and bone mass becomes dysregulated. One such disease, osteoporosis, is characterized by the rapid loss of bone mass and increased risk of fracture. It affects over 200 million people worldwide, with 1 in 3 women and 1 in 5 men projected to have a bone fragility fracture in their lifetimes(*6, 7*). Consequently, understanding cellular sensing mechanisms and the systems-level components that regulate bone mechanotransduction is an area of critical study.

A key biological player in bone mechanotransduction is the osteocyte. These terminally differentiated bone cells express proteins responsible for regulating the behavior of bone-resorbing osteoclasts and bone-building osteoblasts(*8*–*10*). They reside encased in bone matrix and are functionally connected by connexin-43 gap junctions near the ends of long dendritic processes, creating a functional syncytium(*11, 12*). Their location throughout bone and ability to control the behavior of tissue-modifying cells make them ideally designed for their role as resident bone mechanosensors. Osteocytes bear an uncanny resemblance to nerve cells. Both are highly dendritic cells that exist in interconnected networks. Like how nerves sense various stimuli (e.g., heat, pain) in different tissues, osteocytes sense force in bone. Osteocyte mechanotransduction is a complex cell signaling event, and unraveling the critical features that serve as control mechanisms is the subject of ongoing investigation.

Calcium (Ca^2+^) signaling is a ubiquitous second messenger system with pronounced biological influence over nearly every aspect of cell function, including protein conformations, signal transduction, excitability, exocytosis, cell motility, mitochondrial integrity, apoptosis, and gene transcription(*13, 14*). In osteocytes, Ca^2+^ signaling is a key second messenger in mechanotransduction pathways. Activation of Ca^2+^ signaling in osteocytes *in vitro* occurs in response to fluid flow shear stress of similar magnitude to that experienced by endothelial cells(*15*–*21*) as first predicted by Weinbaum and Cowin(*22*). Osteocytes also exhibit increases in cytoplasmic Ca^2+^ in response to substrate deformation(*23*–*26*) and hydrostatic pressure(*27*– *29*). Our group has shown osteocytes use Ca^2+^ signaling to biologically encode tissue level force magnitude *in vivo*(*30*). The variety of components and downstream effects of Ca^2+^ signaling support the theory that it is an important control mechanism for bone mechanoregulation. As such, the mechanisms that regulate Ca^2+^ signaling dynamics in osteocytes represent important features in bone mechanotransduction. One such mechanism recently implicated is acetylcholine receptors.

Acetylcholine (ACh) is a neurotransmitter associated with Ca^2+^ signaling channels that has recently been connected to bone mechanobiology. Neurological disorders like Alzheimer’s disease are correlated with the dysregulation of cholinergic activity. Interestingly, lower bone mineral density(*31*), higher fracture risk(*32*), and osteoporosis(*33*) are all linked to higher rates of dementia. Alzheimer’s patients treated with acetylcholine esterase (AChE) inhibitors, which increase ACh half-life, have the added benefit of 20-30% reduction in hip fracture, improved hip fracture healing, and improved bone quality(*34, 35*). In mice, AChE inhibitors increase trabecular bone volume and reduce bone resorption(*36*). Parasympathetic(*36*) and sympathetic derived ACh impact skeletal mass and development(*37*). All three major bone cells (i.e. osteocytes, osteoblasts, and osteoclasts) express proteins associated with ACh metabolism(*36*–*39*). Despite convincing evidence of the impactful role of ACh in bone biology, the details regarding cholinergic signaling in bone mechanotransduction remain largely unexplored. What’s more, there are very few data describing ACh signaling in osteocyte mechanotransduction. This is largely due to the technical challenges associated with studying osteocytes embedded in mineralized matrix. Our group specializes in studying osteocyte cell biology *in vivo* and has worked to shed light on this subject in the work presented here.

Nicotinic ACh receptors (nAChRs) are ACh-gated ion channels that have high permeability to Ca^2+^(*40, 41*). *Chrna1* encodes for nAChR subunit α1 of the nicotinic receptor, which binds ACh and regulates cellular activation through direct modulation of membrane potential(*42, 43*). Interestingly, the differentiation of osteoblasts into osteocytes is associated with the upregulation of several neurotransmitter-associated genes, including *Chrna1*(*38, 44*). In muscle (i.e., neuromuscular junctions on mature neuromuscular synapses), nAChR α1 subunits are involved in binding ACh and channel gating to control contraction through Ca^2+^ signaling(*45, 46*). In rat bone, *Chrna1* is downregulated after mechanical loading, indicating potential mechanical regulation of nAChRs in bone cells(*44*). As such, ACh may have an influence over osteocyte Ca^2+^ dynamics at the cellular level, and consequently bone mechanoadaptation at the tissue level. Rapsyn, also expressed by osteocytes, is an intracellular scaffolding protein that tightly regulates nAChR mobility on the membrane and clusters nAChRs into functional post-synaptic groups, encoded by the gene *Rapsn*(*47, 48*). This cytosolic protein is critical for proper ACh signaling and is highly expressed in osteocytes (*38*), however very little is currently known about its role in osteocyte function. It is unclear whether ACh-induced tissue-level bone architecture changes in bone result from osteocyte signaling or are mediated through some other mechanism. As such, there is a need for a deeper analysis of the impacts of altered cholinergic signaling in osteocytes on bone mechanotransduction.

We believe the osteocyte serves as a functional neurotransmitter signaling target of the central nervous system in bone. The goal of this study was to determine if cholinergic signaling components nAChR subunit α1 and rapsyn in osteocytes are relevant for bone development and mechanoadaptation. We hypothesized that disrupted osteocyte cholinergic signaling negatively impacts bone tissue organization and responses to mechanical loading. To investigate this question, we generated osteocyte-targeted conditional knockouts (cKOs) for either *Chrna1* or *Rapsn* in mice. We collected skeletal morphometric measurements throughout post-natal development and after anabolic mechanical loading *in vivo*. We also performed an *in vivo* live cell functional assay to determine changes to osteocyte mechanosensitivity in their native environment, with nervous system interactions intact. Our results showed sexually dimorphic differences in bone structure and formation rates between Cre-negative controls and cKO mice. In females, reductions to bone geometry were rescued with anabolic loading, but not in males. Interestingly, female *Chrna1* cKO osteocytes were less mechanosensitive at lower forces, perhaps providing a mechanistic explanation for our other results. Our findings provide strong support the role of osteocytes as direct cholinergic targets impacting bone mechanoadaptation and may assist in defining novel therapeutic targets for bone metabolic diseases (e.g., osteoporosis).

## Results

### Cholinergic nerve fibers are proximate to cortical bone osteocytes *in situ*

Work from others has reported the presence of cholinergic fibers in bone using histological sections, with a particular emphasis on bone directly beneath articular cartilage at the ends of bones(*36*–*39*). Our group has developed an approach for whole mount 3-dimensional imaging of mouse bones to assess the presence of cholinergic nerve fibers. We used a modified BABB (Benzoic Acid Benzyl Benzoate) clearing approach(*49, 50*) to achieve excellent clarity and image quality in intact tissues compared to previous protocols(*50, 51*). We combined this clearing approach with bulk immunolabeling of cholinergic fibers (green) and adrenergic fibers (red) in mouse third metatarsals (MT3s). The cleared and labeled bones were imaged via lightsheet microscopy. In a representative image, it can be seen that cholinergic fibers penetrated the MT3 throughout the length of the bone, with numerous transcortical events at the midshaft (**Fig 1**). An extensive network of labeled structures can be seen in the marrow space. Adrenergic fibers can also be seen with density and arrangement at the ends of the bones that replicate data reported by other groups(*36, 37, 52*). These tissue level imaging data provide strong evidence for the presence of cholinergic nerve fibers in cortical bone and, thus, supports the possibility for functional and/or direct contact between cholinergic nerve fibers and osteocytes.

**Figure 1:**
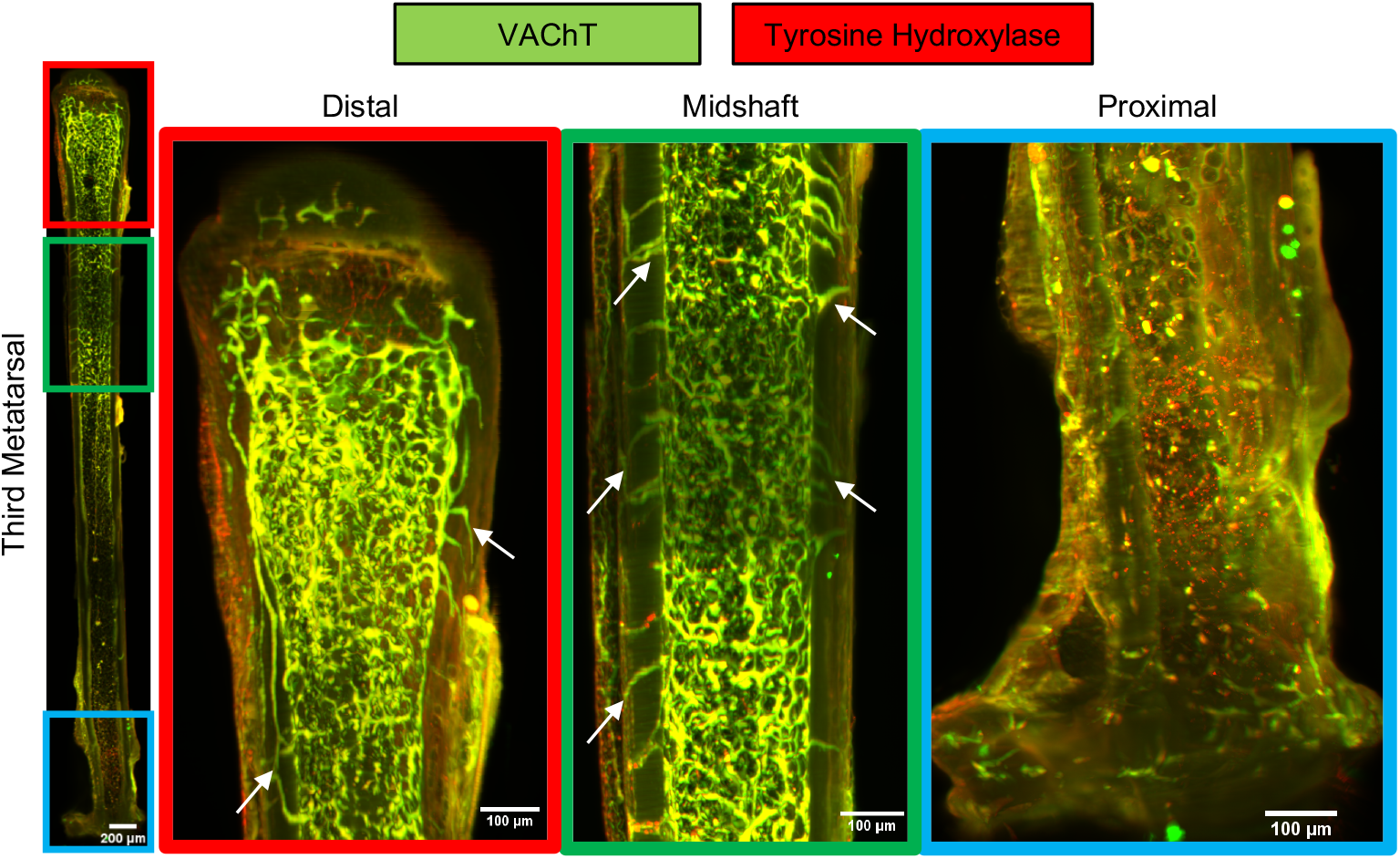
Fluorescent lightsheet imaging of a cleared mouse third metatarsal. Representative 3D projection light sheet microscopy images of a cleared bulk immunohistochemistry-stained mouse third metatarsal with VAChT (Green) and tyrosine hydroxylase (Red) representing cholinergic and adrenergic nerve fibers, respectively. Clear, distinct nerve labeling can be seen at the distal and proximal ends of the bone. At the midshaft, transcortical innervation can be observed, with extensive branching throughout the distal half of the bone. White arrows indicate transcortical cholinergic fibers.

We used an intravital imaging approach to interrogate the prospect of direct contact between cholinergic nerve fibers and osteocytes. Mice expressing GCaMP6f in a CHaT-dependent manner (Strain# 028865, 006410; Jackson Laboratory) were injected with Cy5 fluorescent nanoparticles that are readily taken up by osteocytes *in vivo***(*53*)**. The nanoparticles were injected subcutaneously at the volar aspect of the hindpaw and incubated for 45 minutes. The bones of anesthetized mice were surgically exposed and imaged using 2-photon microscopy. Very fine green cholinergic nerve fibers were seen throughout the cortex (**Fig 2A**). Interestingly, these fibers occasionally interfaced directly with red-labeled osteocytes, as indicated by yellow overlap signal (**Fig 2B-C**). When the representative z-stack is rotated 90-degrees, the interaction between some osteocytes and nerves becomes readily apparent (**Fig. S2**). We evaluated the the distribution of osteocytes within 100μm of cholinergic nerve fibers in metatarsal cortical bone in mice *in vivo* (**Fig 2D-F**). Female mice exhibited a more skewed distribution compared to males (**Fig. 2E-F, Fig. S3**). These data validate the presence of cholinergic nerves in the osteocyte microenvironment and strongly suggest direct interaction between nerves and osteocytes. Moreover, they show that embedded nerve fibers are present within mineralized matrix tissue in cortical bone.

**Figure 2:**
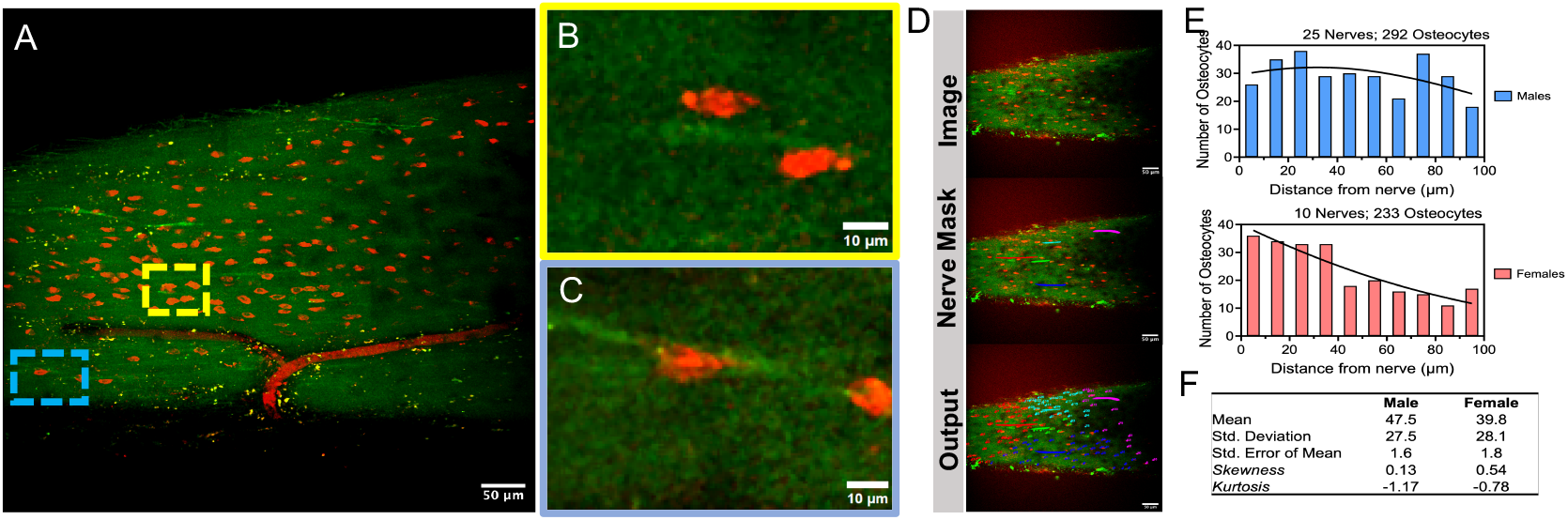
Osteocytes are positioned close to cholinergic nerves in vivo. Intravital Z-stack image (40μm thick) of metatarsal bone osteocytes in mice expressing GCaMP6f in acetylcholine transferase expressing cells taken using two photon microscopy at about 20μm below the periosteal surface (A). The red fluorescence signal is driven by a 15mL volume of ultrasmall RGD-functionalized integrin targeting nanoparticles (5-7nm) at 10μM concentration. This static 3D image indicates the colocalization of osteocytes, labeled in red, with cholinergic nerve fibers, labeled in green. (B-C) Yellow and Blue insets depict single 2D frames where the nerve can be seen directly interacting with osteocytes. (D) Representative images for nerve-osteocyte proximity analysis showing the input image with osteocytes in red and ChAT signal containing cholinergic nerve fibers in green. The masks were drawn over the nerve fibers of the input image, and proximity analysis of the shortest distance between each osteocyte and a nerve was calculated and represented in the output image. (E) Histogram with an overlaying Gaussian curve showing the distances of osteocyte distribution around the nerve in both sexes. (F)Both males and females fail the normality test (p value<0.0001), and a significant difference (p value=0.001) exists in the distribution of osteocytes between the sexes (Fig S3). Kolmogorov-Smirnov test was used for normality assessment. Mann-Whitney non-parametric test was used for comparing between sexes.

### Osteocyte-targeted Deletion of ACh Receptor Components Alters Load-Induced Ca^2+^ Dynamics

We observed osteocyte Ca^2+^ signaling responses to mechanical loading *in vivo* to test the functional impact of cholinergic signaling on osteocyte mechanotransduction. Osteocyte-targeted GCaMP6f-expressing mice were generated with concurrent conditional deletion (cKO) of *Chrna1* or *Rapsn*. Osteocytes were imaged intravitally for each mouse group at the dorsal mid-diaphysis of the MT3 during active mechanical loading. We investigated the strain dose response by subjecting bones to strain values between 250-3000με. For all groups, the percentage of responding osteocytes as indicated by Ca^2+^ signaling increased with increasing strain levels as expected based on previous studies (**Fig 3**)**(*30*)**. In female mice, *Chrna1* cKO mice had fewer responsive osteocytes at 250 and 500με loading. Additionally, *Rapsn* cKO mice had a lower number of responding cells at 250με (**Fig 3C**). These data indicate a decreased mechanosensitivity among osteocytes at lower force levels *in vivo*. There were no differences to the change in fluorescent intensity for the responding cells in female mice.

**Figure 3:**
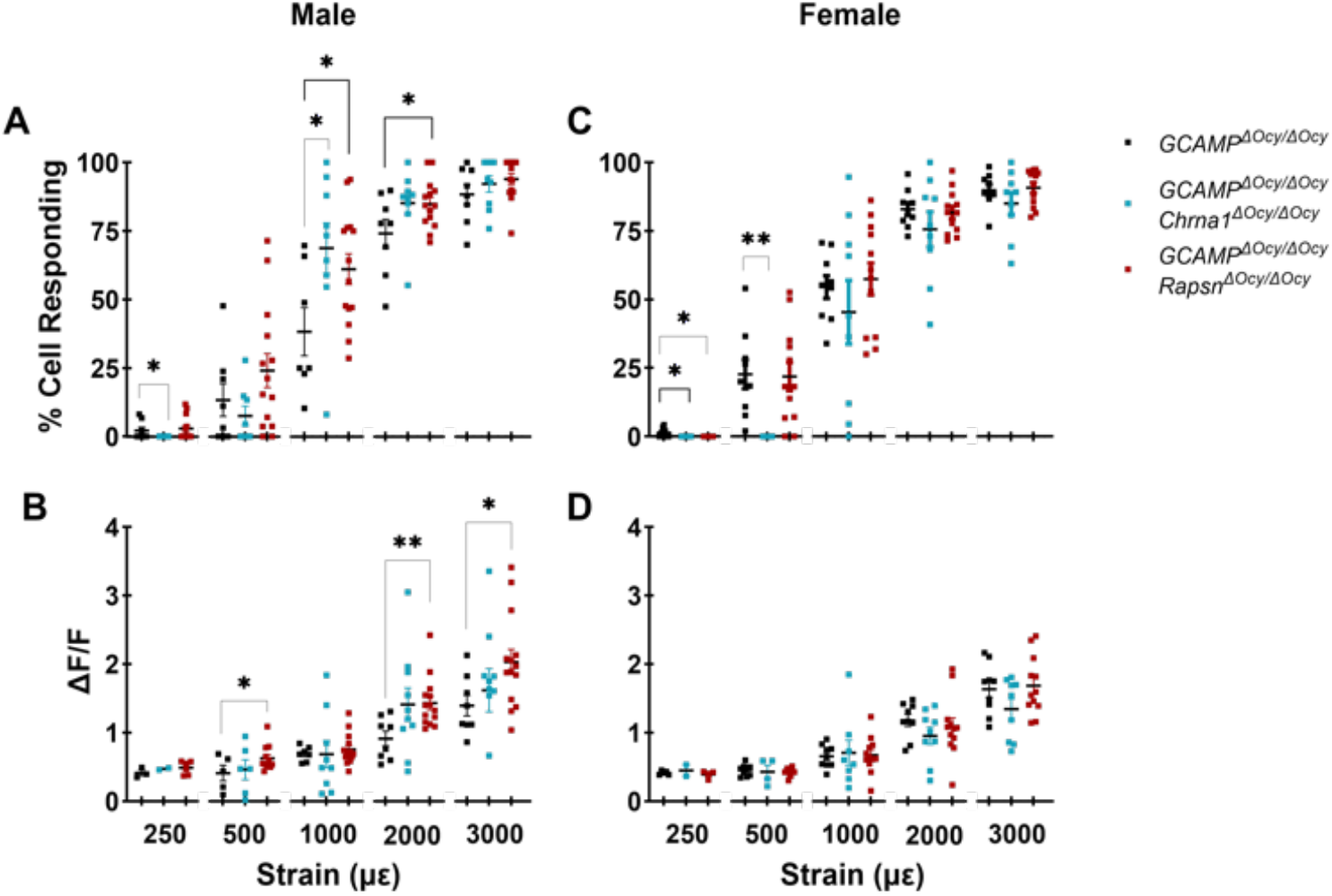
*In vivo* osteocyte mechanotransduction is impacted by *Chrna1* and *Rapsn* cKO in a sexually dimorphic way. Intravital imaging of osteocyte Ca^2+^ signaling responses to mechanical loading was used to assess molecular level functional impacts of either *Chrna1* or *Rapsn* cKO. (A) In males, the percentage of responding osteocytes decreased with Chrna1 cKO at very low strains but increased at higher strains for both *Chrna1* and *Rapsn* cKO. (B) Interestingly, *Rapsn* cKO also increased response intensity among responding cells at 3 of the 5 strain levels tested. These results indicate dysregulation of mechanosensation in male osteocytes with ACh signaling disruption. (C-D) While female response intensities were unaltered, the number of responding cells was decreased at lower strain values tested for *Chrna1* cKO mice compared to controls. T-tests were performed across groups within each strain level (*p>0.05; **p>0.01). Sample sizes are as follows: % Cells Responding: [GCaMP: 7-9, *Chrna1*: 9-10, *Rapsn*: 12-14)]. Delta F: [GCaMP: 3-9, *Chrna1*: 2-10, *Rapsn*: 4-14)].

The results in male mice were categorically different. The number of responsive cells at lower magnitude values was largely unchanged, except for a slight but significant reduction in *Chrna1* cKO mice at 250με compared to controls. At 1000 and 2000με *Rapsn* cKO males had a higher number of responsive osteocytes compared to controls (**Fig 3A**). Likewise, *Chrna1* deletion led to an increase in responding cells at 1000με (**Fig 3A**). Moreover, the intensity among responding cells was increased at 500, 2000, and 3000με for *Rapsn* cKO mice (**Fig 3B**). These results indicate a general trend for increased mechanosensitivity in male mice with constitutively dysfunctional ACh signaling. Interestingly, this trend appears to be centered on the mid-high range of physiological loading magnitudes.

Endogenously expressed GCaMP6f signal in osteocytes was used to evaluate 2D cellular morphology *in vivo*. There were no differences in the number of cells between any of the groups (**Fig. 4A**). Female *Chrna1* cKO osteocytes had a slightly higher cell area when compared to GCaMPf controls (**Fig. 4B**). For both sexes, *Chrna1* cKO cells were more elongated and less round compared to controls (**Fig. 4C-F**). *Rapsn* cKO did not alter 2D osteocyte morphology in females (**Fig. 4A-F**). In *Rapsn* cKO males, again cells were more elongated (**Fig. 4H,J,K**). These results suggest that Ach signaling disruption results in altered osteocyte shape, which could explain the impacts on *in vivo* osteocyte mechanotransduction.

**Figure 4:**
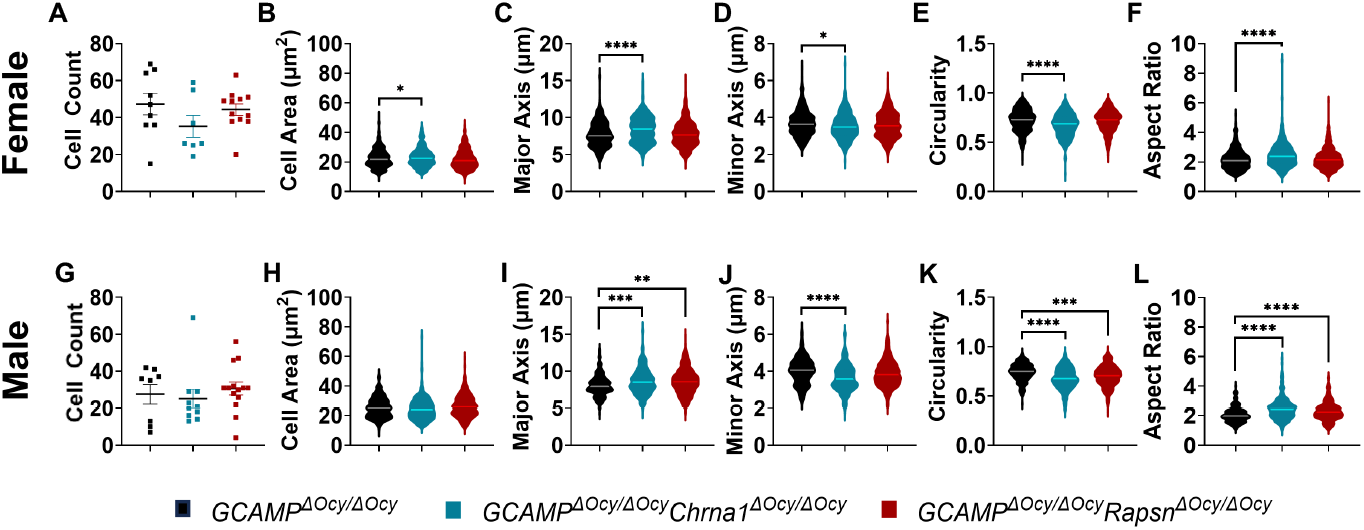
*In vivo* osteocyte morphology is impacted by ACh signaling disruption in a sexually dimorphic way. Endogenously expressed GCaMP6f signal in osteocytes was used to evaluate 2D cellular morphology *in vivo*. (A) There were no differences in the number of cells per area between the groups. (B) Female *Chrna1* cKO osteocytes have a slightly lower cell area when compared to GCaMP controls. For both females (C-F) and males (I-L), *Chrna1* cKO cells were more elongated and less round than controls. (A-F) *Rapsn* cKO did not alter 2D osteocyte morphology in females. In *Rapsn* cKO males, cells were again more elongated (J-L). These results suggest that ACh signaling disruption produces altered osteocyte shape, which could explain the impacts on *in vivo* osteocyte mechanotransduction. Two-way ANOVA with Tukey’s multiple comparison test was performed with cKO and GCaMP controls to determine significance (*p<0.05; **p<0.01, ***p<0.001, ****p<0.0001) for each metric. N = 4 - 69 cells per mouse, group sample size [GCaMP: 7-9, *Chrna1*: 9-10, *Rapsn*: 12-14].

### Osteocyte-targeted *Chrna1* conditional deletion leads to site-specific skeletal abnormalities

The α1 nicotinic subunit is responsible for ACh binding during receptor activation. To assess the impact of α1 nicotinic cholinergic receptor function in osteocytes, we achieved osteocyte-targeted cKO of *Chrna1* using a 10kb DMP-1 targeted Cre-Lox system (**Fig. S1**). DEXA scans of the whole body, hind limbs, and lumbar spine were performed at intermittent periods between the ages of 4 and 21 weeks. Comparisons were made between *Chrna1*^Δ*Ocy/*Δ*Ocy*^ mice and littermate aged matched *Chrna1*^+*/*+^ controls. We found that *Chrna1*^Δ*Ocy/*Δ*Ocy*^ mice generally formed skeletons with normal bone mineral density (BMD) compared to *Chrna1*^+*/*+^ controls (**Fig 5A-F**). The only exception was in older female mice, where cKO mice had increased BMD in the lumbar spine at 17 weeks (+10.6%; **Fig 5B**) and 21 weeks (+11.7%; **Fig 5B**). µCT analysis of adult tibiae indicated sexually dimorphic site-specific differences in tissue development (**Fig 6A-J** Non-loaded, **Fig. S4**). *Chrna1*^Δ*Ocy/*Δ*Ocy*^ females exhibited reduced bone mass at the metaphysis compared to *Chrna1*^+*/*+^ mice. This included reductions in trabecular bone volume over total volume (−32.7%; **Fig 6B**), the number of trabeculae (−10.6%; **Fig 6C**), and the density of total volume (−20.4%; **Fig 6D**). By contrast, *Chrna1*^Δ*Ocy/*Δ*Ocy*^ males exhibited a midshaft phenotype with reduced cortical bone parameters. They had midshafts with lower mean total area (−15.2%; **Fig 6G**), resulting in lower minimum moment of inertia (−33.6%; **Fig 6H**), lower maximum moment of inertia (−29.7%; **Fig 6I**) and lower polar moment of inertia compared to *Chrna1*^+*/*+^ males (−31.1%; **Fig 6J**). These results collectively indicate that disrupting cholinergic signaling in osteocytes using constitutive deletion of *Chrna1* or *Rapsn* leads to slight sexually dimorphic differences in skeletal development (**Tables S1-2**). While the results are not uniform between the sexes, there is a clear role for osteocyte ACh signaling in bone development.

**Figure 5:**
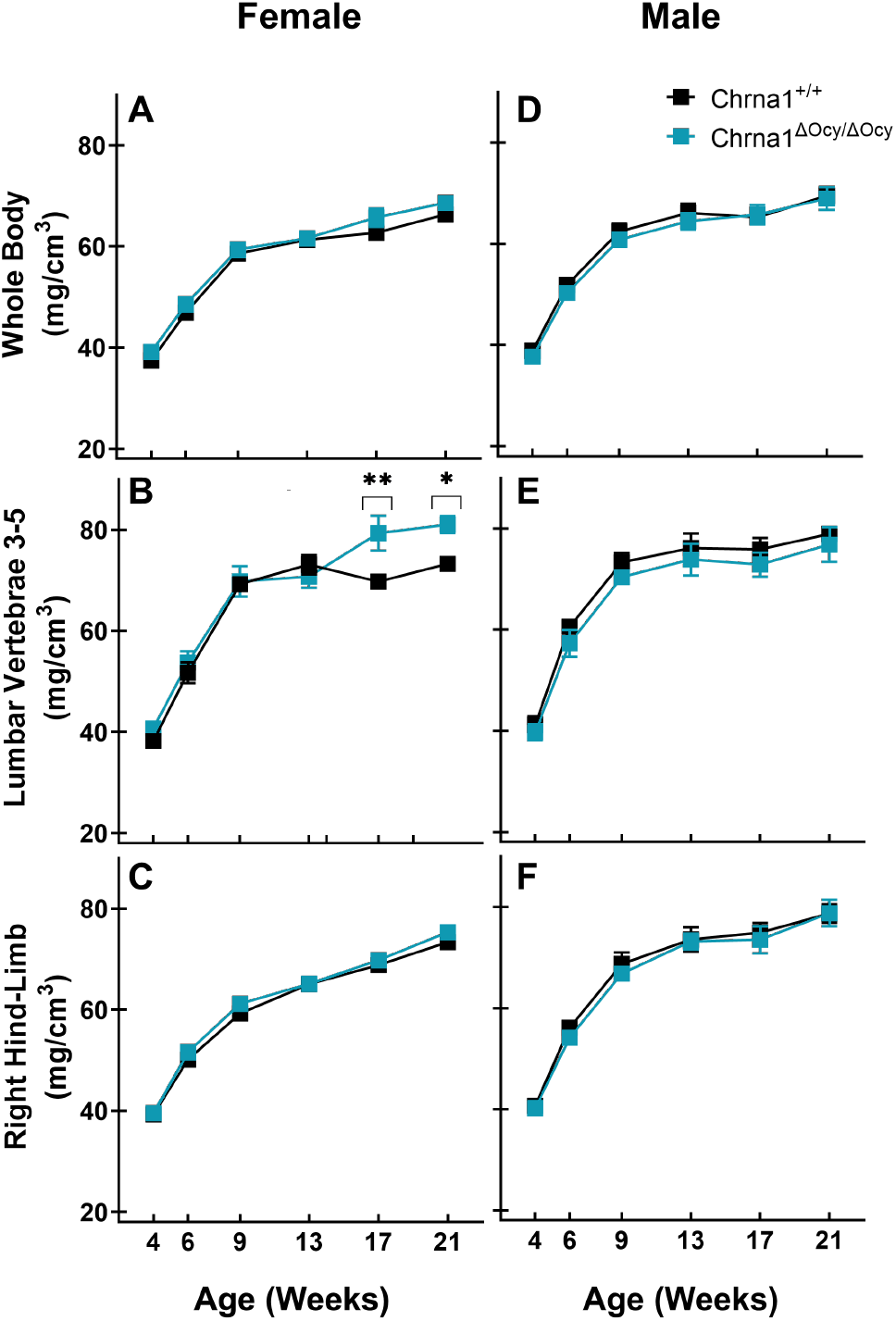
*Chrna1* cKO produces mostly normal skeletons during development. Bone mineral content and density were measured using DEXA for the whole body, lumbar spine, and right hind limb between 4 and 21 weeks of age. We found no differences between control and *Chrna1* cKO groups in either female (A-C) or male (D-F) mice. Two-way ANOVA was performed with interactions between genotype and age. N = 6-15/group. A post-hoc Bonferroni test was computed across genotypes for each timepoint to determine significance (*p<0.05; **p<0.01). N = 6-13 per group and skeletal location.

**Figure 6:**
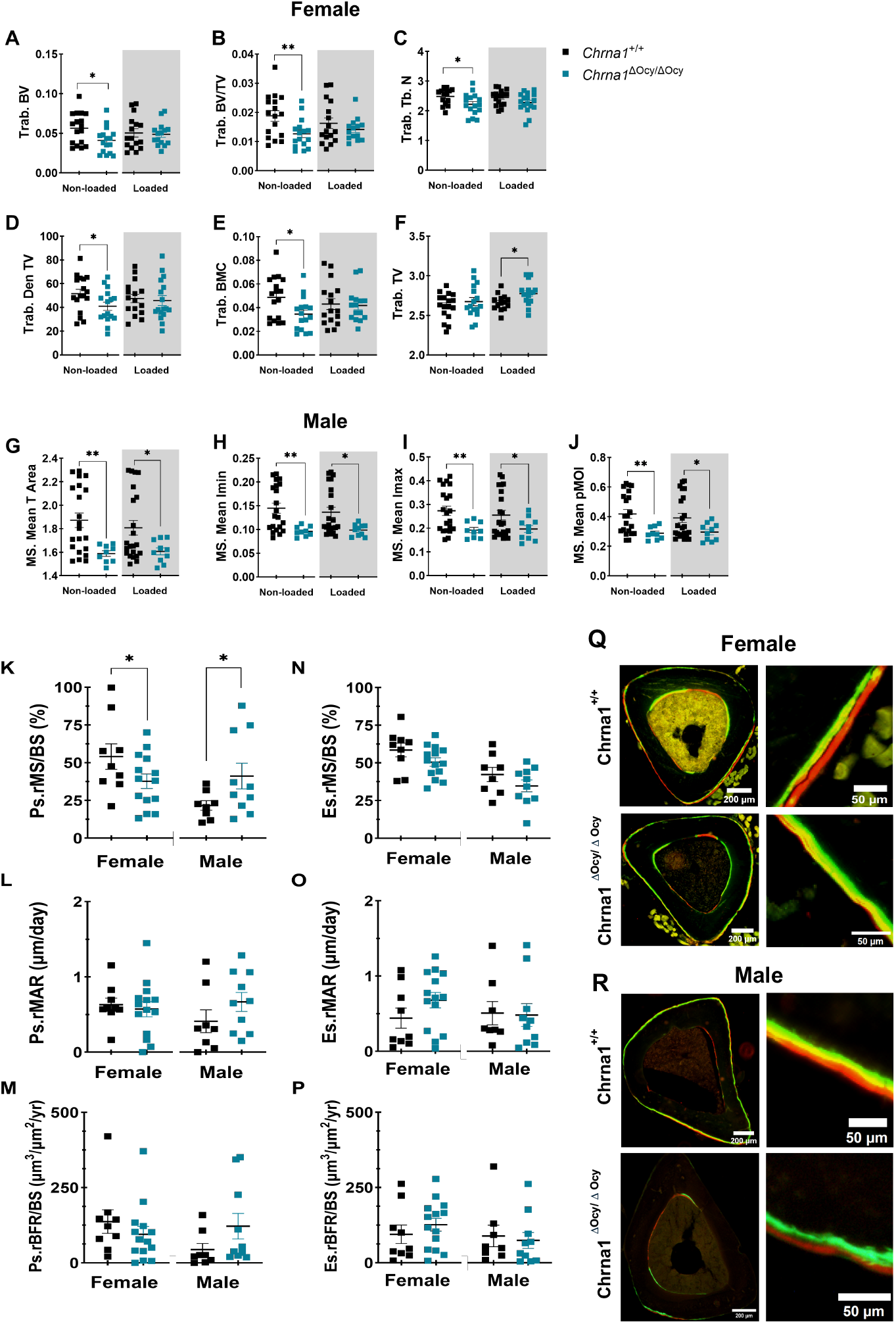
*Chrna1* cKO impaired the anabolic response to mechanical loading. Adult mice were subjected to tibial loading, and bones were excised 12 days later. µCT measured tibial geometry for females (A-F) and males (G-J) to compare non-loaded limbs (white background) with loaded limbs (shaded grey background). Female Chrna1 cKO mice decreased trabecular bone measures(B-D) at baseline. Likewise, male *Chrna1* cKO mice had a decreased mid-shaft bone phenotype (H-J). Interestingly, female trabecular bone was recovered with loading (B-D). There was no change in male mid-shaft metrics with loading (H-J). (K-P) Fluorochrome labeling with calcein followed by alizarin red was used for osteoblast-derived indices of dynamic histomorphometry. Relative metrics were computed by subtracting non-loaded limbs from loaded limbs. Indices were calculated for both the endosteal and periosteal surfaces. Very few load-induced formation differences were measured between the control and cKO groups. Periosteal mineralizing surface was decreased in female cKO mice and increased in male cKO mice (K). Student T-tests were performed between cKOs and Cre-negative controls to determine significance (p<0.05). (p<0.05). N= 8-22 per group and assay.

### Osteocyte-targeted *Chrna1* Deletion Impairs Bone Anabolic Response to Mechanical Challenge

To assess the impact of osteocyte cholinergic signaling on bone mechanotransduction, we studied tissue changes in response to *in vivo* loading over time. Mice received *in vivo* tibial loading to peak calibrated strain magnitudes of −2250µε 3 times per week for 2 weeks. Post-mortem analyses of loaded and non-loaded tibia included µCT and dynamic histomorphometry. The anabolic impact of mechanical loading was present but mild in control mice (**Fig. S4**). As noted above, non-loaded baseline conditions in tibiae from *Chrna1*^Δ*Ocy/*Δ*Ocy*^ females exhibited reduced trabecular bone mass at the metaphysis compared to Cre-negative littermate controls. This phenotype was rescued in loaded bones (**Fig 6A-E** Loaded), with trabecular total volume (Trab. TV; **Fig 6F** Loaded) even exceeding that of *Chrna*^+*/*+^ controls by +4.2%. Non-loaded tibiae from *Chrna1*^Δ*Ocy/*Δ*Ocy*^ males had normal trabecular bone but exhibited a midshaft phenotype with reduced cortical bone area and moment of inertia. Unlike females, anabolic loading did not rescue the phenotype in male mice(**Fig 6G-J**, grey background). Relative dynamic histomorphometry indices were assessed by subtracting non-loaded values from loaded limbs. *Chrna1* cKO led to a slight decrease in periosteal formation rates compared to controls in female mice. By contrast, this measure was increased in male cKO mice (**Fig 6K**). No other differences were found in load induced formation markers for either sex (**Fig 6K-P**).

### Osteocyte-targeted *Chrna1* Deletion Impacts Toughness in Male Femurs

We next set out to determine if there would be changes to the whole bone mechanical properties at skeletally maturity. Femurs from 16-18-week mice underwent monotonic three-point bending to failure. Female *Chrna1*^Δ*Ocy/*Δ*Ocy*^ mice were not different than Cre-negative littermate controls (**Fig 7A-D**). Conversely, male *Chrna1*^Δ*Ocy/*Δ*Ocy*^ mice had greater plastic deformation, indicating their bones were more ductile and resistant to mechanical failure. Male *Chrna1*^Δ*Ocy/*Δ*Ocy*^ mice had 228% increased post-yield displacement (**Fig 7F**), 29.1% reduced fracture force (**Fig 7G**), and +98.7% increased failure energy (**Fig 7H**) compared to Chrna1^+*/*+^ controls. FTIR tissue composition analysis found no changes to the mineral-to-matrix ratio, collagen maturity, or mineral crystallinity for cKO compared to Cre-negative controls for either sex (**Fig 7I-J**). Together, these results firmly support a role for osteocyte ACh signaling in formation and/or regulation of bone material properties in male mice.

**Figure 7:**
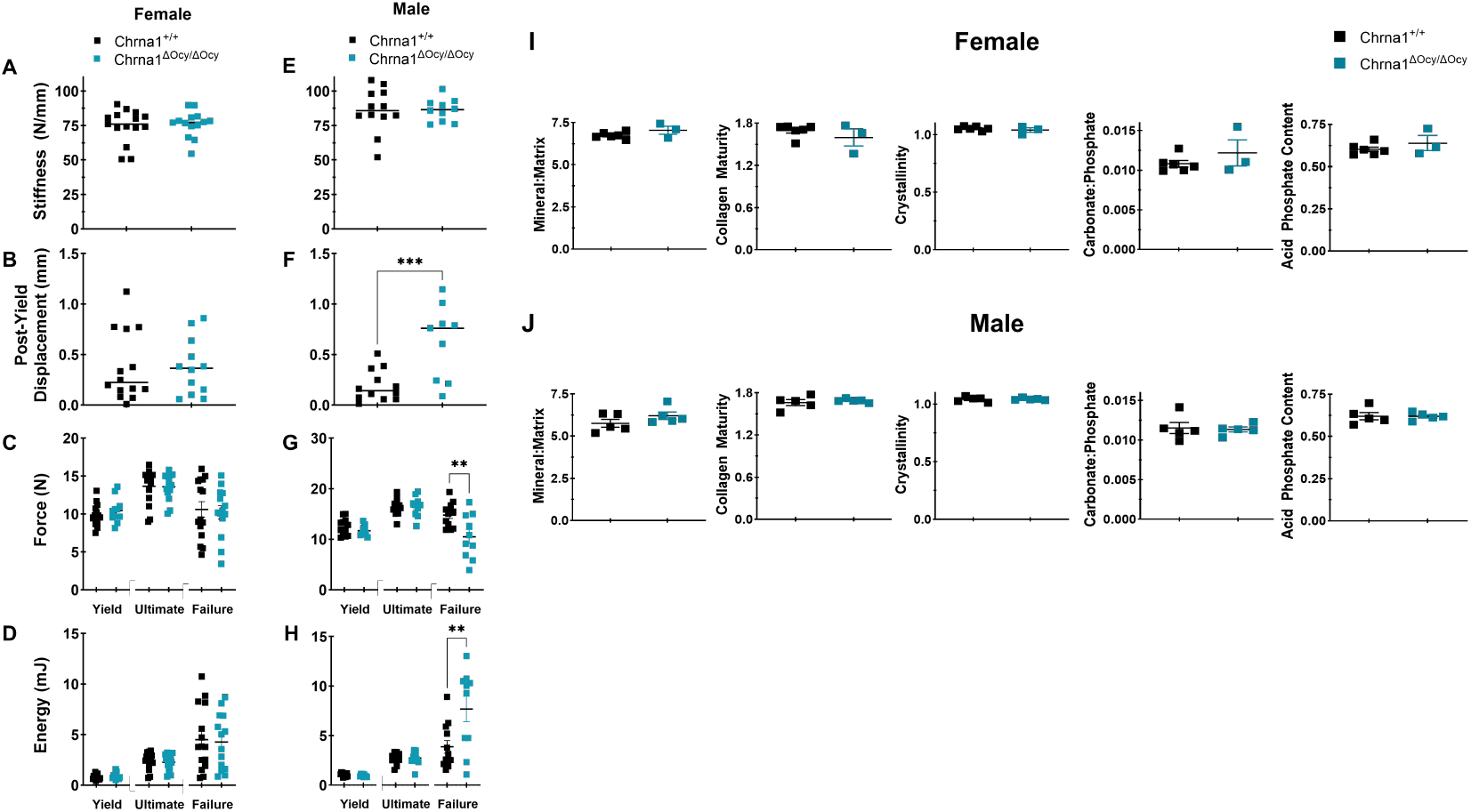
*Chrna1* cKO increased femur ductility and toughness in males. (A-D) Mechanical properties were assessed with load-displacement curves from monotonic loading to failure. There were no differences in female bones. (E-H) Male *Chrna* cKO mice had increased post-yield deflection and total work to failure, indicating more ductile tissue properties. FTIR was used to evaluate bone compositional properties from a subset of left femora from mice used for mechanical testing for females (I) and males (J). No differences were found between the control and cKO groups. Student T-tests were performed with cKO and Cre-negative controls to determine significance (p<0.05) for each metric. N = 3-15 per group and assay.

### Osteocyte-targeted *Rapsn* cKO Leads to Site-Specific Skeletal Abnormalities

Rapsyn clusters cholinergic ion channels on the cell membrane, allowing for coordinated receptor/channel function and more potent responses. We repeated the same experiments performed above with *Chrna1* cKOs in mice lacking *Rapsn* in osteocytes (**Fig S1**). To study growth and development, mice with *Rapsn* cKO had DEXA scans of the whole body, hind limbs, and lumbar spine over time. We found that cKO mice of both sexes largely formed skeletons with normal BMD compared to Cre-negative littermate controls (**Fig 8A-F**). Males exhibited a spike in BMD at 6 weeks in the lumbar spine that resolved and normalized in subsequent weeks (**Fig 8E**). We also evaluated tibia morphology using µCT analysis (**Tables S3-4**). We observed decreases in bone volume indices in female bone; including cortical volume (−5.1%; **Fig 9A**), cortical total volume (−8.3%; **Fig 9B**), the total volume (−8.3%; **Fig 9E**). *Rapsn* cKO mice had increased bone mineral density compared to controls (+1.3%; **Fig 9D**). Male *Rapsn*^Δ*Ocy/*Δ*Ocy*^ mice had increased bone volume density at the midshaft (+1.2%; **Fig 9H**) compared to Cre-negative littermate controls but exhibited no other differences. Again, we see here sexually dimorphic impacts of ACh signaling disruption in osteocytes (**Table S8-10**).

**Figure 8:**
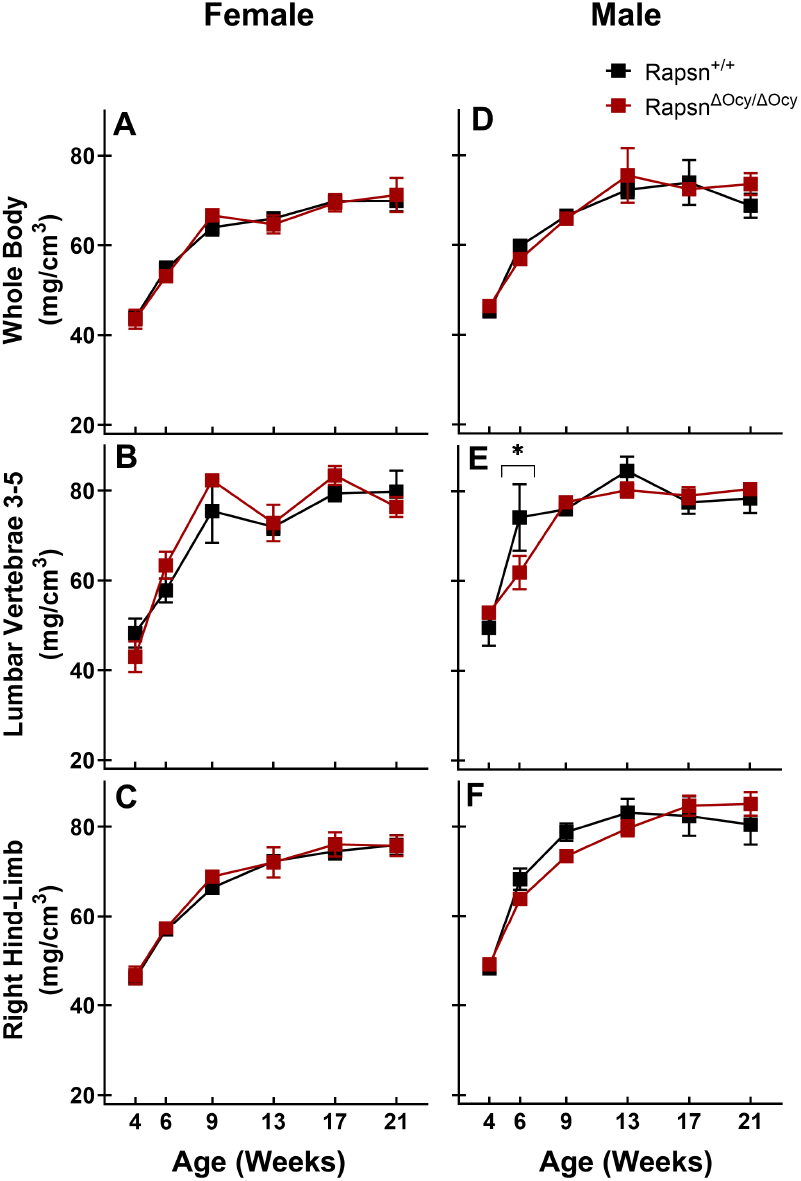
*Rapsn* cKO produces normal skeletons during development. Bone mineral content and density were measured using DEXA for the whole body, lumbar spine, and right hind limb between 4 and 21 weeks of age. We found no differences between control and *Chrna1* cKO groups in either female (A-C) or male (D-F) mice. Two-way ANOVA was performed with interactions between Genotype and Age. N = 3-7/group. A post-hoc Bonferroni test was computed across genotypes for each timepoint to determine significance (*p<0.05).

**Figure 9:**
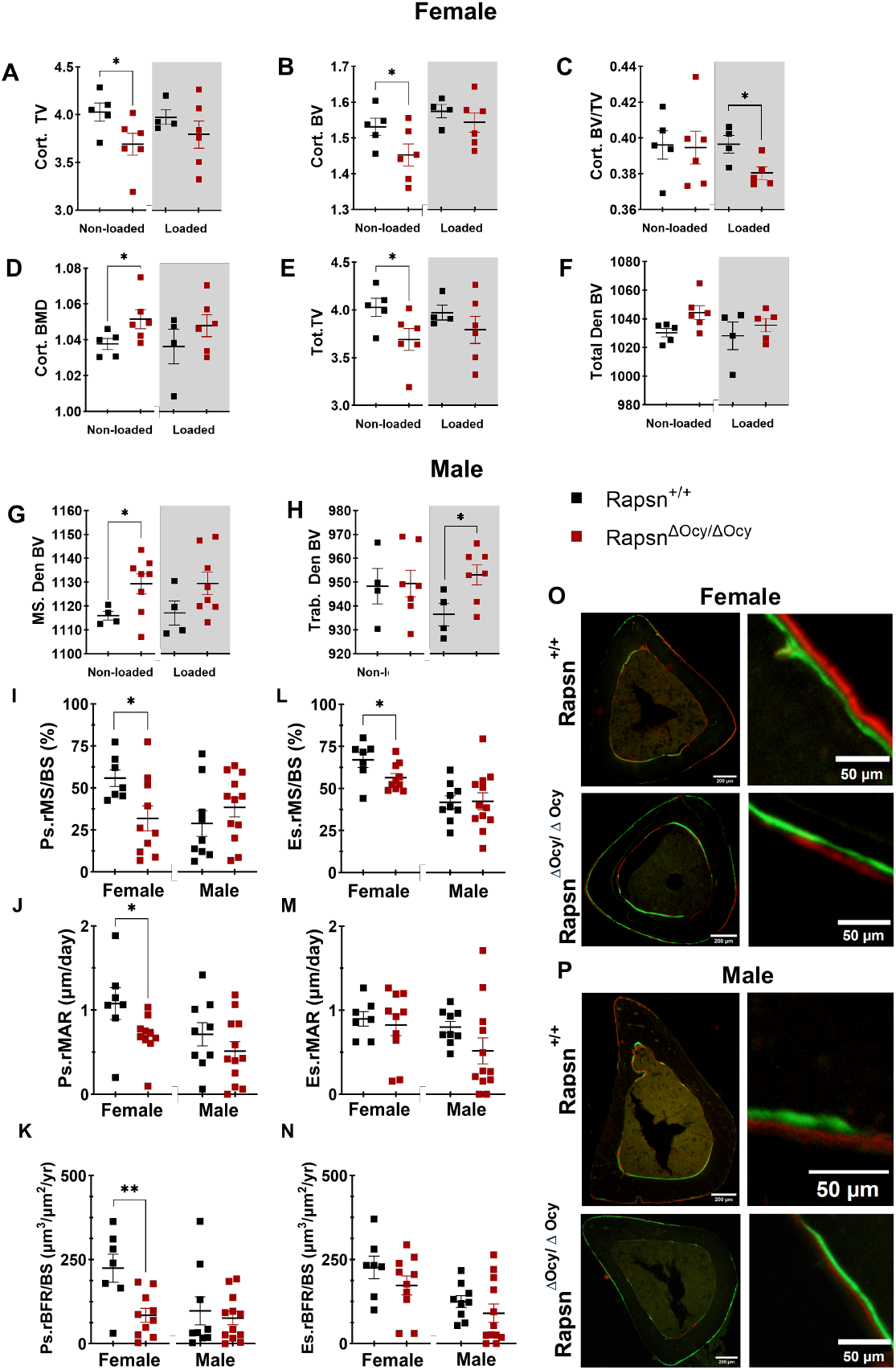
*Rapsn* cKO impaired bone formation in response to mechanical load. Adult mice were subjected to tibial loading, and bones were excised 12 days later. µCT measured tibial geometry for females (A-F) and males (G-H) to compare non-loaded limbs (white background) with loaded limbs (shaded grey background). Female Rapsn cKOs mice at baseline had lower overall volume indices (A-B) but increased bone mineral density (D). These results imply increased tissue hardness to compensate for lower volume/mass. These values are normalized with anabolic mechanical loading. (G) Male *Rapsn* cKO mice had increased tissue density at baseline compared to controls. (H) This difference resolved with loading, while trabecular tissue density increased. (I-P) Fluorochrome labeling was used for osteoblast-derived indices of dynamic histomorphometry. Relative metrics were computed by subtracting non-loaded limbs from loaded limbs. Indices were calculated for both the endosteal and periosteal surfaces. Scale bars are 200μm or 50μm for inset. Female *Rapsn* cKOs exhibited reduced bone formation rates (I-L), suggesting decreased mechanosensitivity. No differences were found between male control and cKO mice. Student T-tests were performed between cKOs and Cre-negative controls to determine significance (p<0.05). N = 4-12 per group and assay.

### Osteocyte-targeted *Rapsn* Deletion Impairs Bone Anabolic Response to Mechanical Challenge

To assess the impact of *Rapsn* on bone mechanotransduction, we studied tissue changes in response to *in vivo* loading. Mice received *in vivo* tibial loading to peak calibrated strain magnitudes of 2250με 3 times per week for 2 weeks. Postmortem analyses of loaded and non-loaded tibia included microCT and dynamic histomorphometry. Non-loaded tibiae from *Rapsn*^Δ*Ocy/*Δ*Ocy*^ females exhibited deterioration in the metaphyseal cortex. Loading rescued this phenotype, returning all deficiencies to control values (**Fig 9A-B, E**, grey background). However, loading was not beneficial overall. *Rapsn* cKO mice suffered decreased bone volume fraction (BV/TV) with anabolic loading compared to control mice (−4.1%; **Fig 9C**). These results indicate an increase in the total volume (or cross-sectional area) of the bone during load-induced adaptation. In male *Rapsn*^Δ*Ocy/*Δ*Ocy*^ mice, loading only caused increased bone volume density in the metaphyseal trabeculae (+1.8%; **Fig 9H**).

Relative dynamic histomorphometry indices were assessed by subtracting non-loaded values from loaded limbs. The anabolic impact of mechanical loading was present but mild in control mice (**Fig. S5**). *Rapsn*^Δ*Ocy/*Δ*Ocy*^ females suffered from lower metabolic rates on the periosteum compared to Cre-negative littermate controls, with suppression of relative mineral apposition rate (Ps.rMAR) by −70.6% (**Fig 9I**), relative mineralizing surface (Ps.rMS/BS) by −25.6% (**Fig 9J**), and relative bone formation rate (Ps.rBFR) b y −88.2% (**Fig 9K**). Females also had reduced endosteal relative bone formation rate (−55.7%; **Fig 9L**). In males, *Rapsn*^Δ*Ocy/*Δ*Ocy*^ mice showed no differences compared to Cre-negative littermate controls.

### Osteocyte-targeted *Rapsn* cKO Maintains or Minimally Increases Whole-Bone Level Mechanical Competence in Femurs

We next set out to determine if there would be changes to the whole bone mechanical properties at skeletally mature bone mass (**Fig 10**). Femurs from 16-18-week mice underwent monotonic three-point bending to failure. Female mice did not show any differences in mechanical properties, whereas *Rapsn*^Δ*Ocy/*Δ*Ocy*^ males exhibited +10.9% higher ultimate force than controls (**Fig 10G**). After measuring the tissue composition of a subset of femurs used for mechanical testing, we determined no changes to the mineral-to-matrix ratio, collagen maturity, or mineral crystallinity for cKO compared to Cre-negative controls (**Fig 10I-J**).

**Figure 10:**
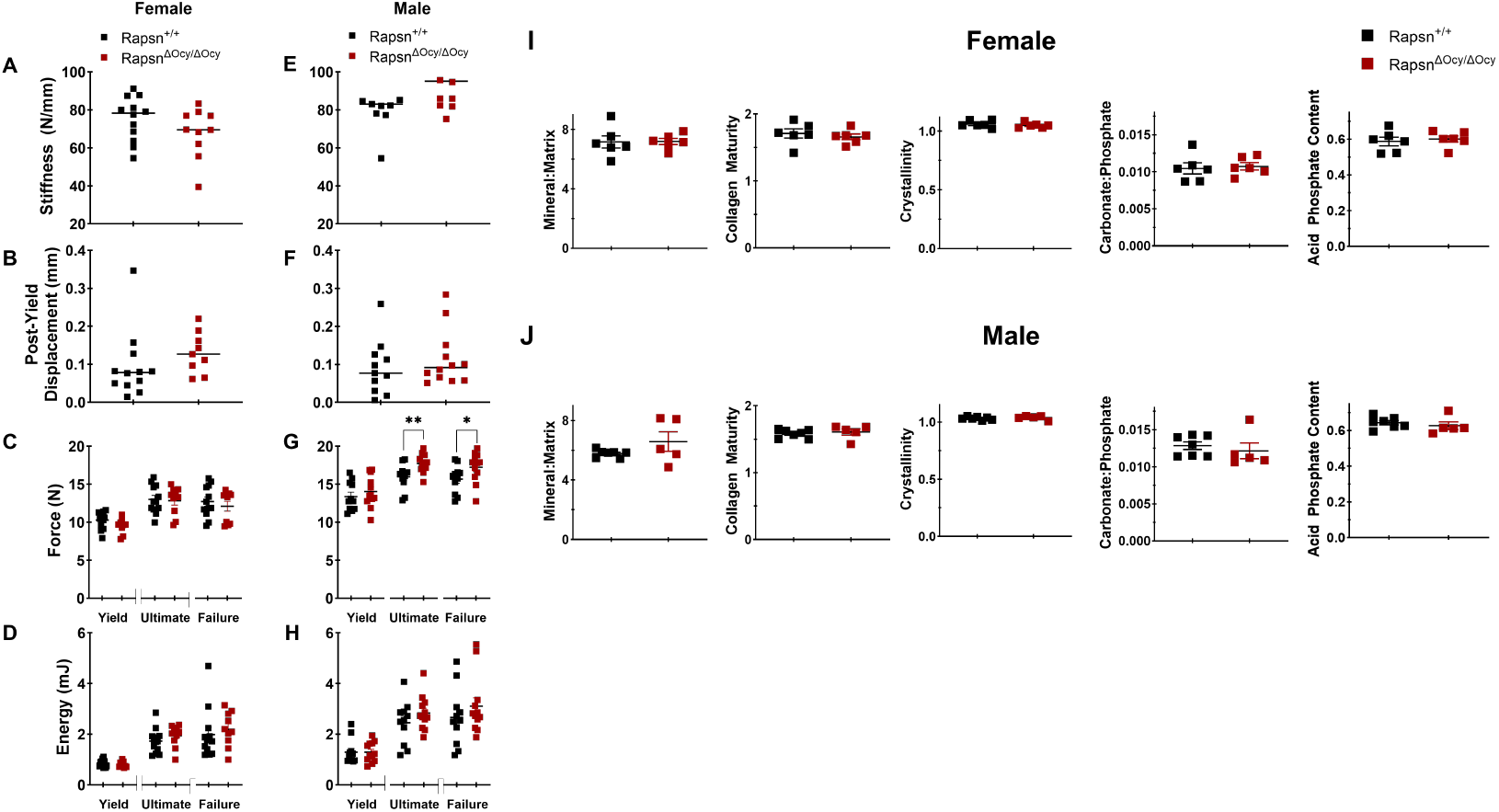
*Rapsn* cKO did not alter whole bone mechanical properties. Mechanical properties were assessed with load-displacement curves from monotonic loading to failure. (A-D) There were no differences in female bones. (E-H) Male *Rapsn* cKO mice had increased ultimate and failure forces, indicating a slight improvement in tissue strength. FTIR was used to evaluate bone compositional properties from a subset of left femora from mice used for mechanical testing for females (I) and males (J). No differences were found between the control and cKO groups. Student T-tests were performed with cKO and Cre-negative controls to determine significance (p<0.05) for each metric. N = 5-12 per group and assay.

## Discussion

Here, we report the first ever evidence of direct physical interaction between embedded osteocytes and cholinergic nerve fibers *in vivo*. These exciting findings imply a functional interaction between the peripheral nervous system and osteocyte networks. Osteocytes use Ca^2+^ signaling to translate tissue level mechanical load magnitude information into biological signals(*30*). nAChRs are Ca^2+^ channels that help to regulate membrane potential in muscles, thus determining one aspect of muscle activation sensitivity(*54, 55*). Given osteocytes are primarily mechanosensors and the role for nAChRs in Ca^2+^ signaling, we suggest a new signaling paradigm wherein cholinergic nerves regulate some aspects of osteocyte mechanosensitivity. This paradigm is supported by studies reporting anabolic impacts of cholinergic therapeutics on fracture risk(*34, 35*) and bone mass(*36, 37*). Moreover, our work is corroborated by work from others showing that osteocytes respond to ACh stimulation(*56*) and express nAChRs genes(*38*) that are differentially expressed after mechanical loading(*44*). Our data, as far as we are aware, are the first to test the impact of osteocyte targeted conditional deletion of ACh receptor components. Additionally, we offer novel intravital imaging approaches linking live osteocyte functional outcomes *in vivo* to their ACh receptor function directly with endocrine and paracrine physiology fully intact. These data work to fill a critical knowledge gap in basic bone biology. For most tissue systems, we understand the homeostatic balance achieved by parasympathetic and sympathetic signaling. In bone, however, much more is known about the role of sympathetic signaling(*57*–*59*) than parasympathetic and/or cholinergic signaling. The data offered here show the first evidence that there is a functional link between osteocytes and cholinergic signaling, suggesting a new physiological link that could be important for bone homeostasis and, thus, novel therapeutic targets.

An important consideration includes the spatial patterning of neurons in the bone and their relation to bone cells. Using a novel tissue-clearing protocol produced by our lab, we successfully cleared whole bones for 3D characterization of skeletal innervation. Cavities that acted as entry points into the marrow space for large nerve fibers were found to penetrate various regions of the cortical shell, with higher densities at the distal and proximal ends compared to the mid-shaft. The embedded position of these nerves suggests opportunity for neurotransmitter communication with osteocytes. Moreover, we showed plausible interactions between cholinergic nerves and osteocytes. These exciting initial insights motivate future studies aimed at defining the details of this physical interaction as well as outlining the nature of the functional behavior at those sites. One meaningful limitation to these data is that only larger positively stained bodies are currently resolved. Future work should include larger load-bearing long bones, such as the femur and tibia.

We achieved conditional osteocyte-targeted deletion of *Chrna1* and *Rapsn*, which caused impairments to bone structure and bone anabolism in a sexually dimorphic manner. Female *Chrna1* cKOs showed deteriorated microarchitecture tibial trabecular bone while female *Rapsn* cKO showed reductions cortical bone. Male *Chrna1* cKO mice suffered from smaller cortical shells at the midshaft in the tibia with increased ductility and toughness in the femurs. Broad phenotypic changes in BMD during skeletal development were not observed. The site and sex specific impact of our cKO models highlights the complexity of the osteocyte-ACh signaling axis and implies redundant regulatory mechanisms. This is common for other key features of bone mechanotransduction like membrane hemichannels(*60, 61*) and integrins(*23, 62*). It is important to note that high resolution microCT data were collected at the end of our experiment from adult skeletally mature mice. AChE levels peak in early development to guide endochondral ossification before drastically declining, maintaining low levels for most of the life(*63*). As such, it is possible that more drastic or different phenotypic changes are present in earlier developmental stages than what we tested here. Further investigation including earlier ages (<4 weeks) is needed to interrogate periods of elevated developmental cholinergic signaling. Likewise, it is important to follow these studies up with data from aging mice. Reduced ACh and increased AChE levels may occur in the elderly and are both associated with age-related diseases such as dementia((*64, 65*). Thus, how cholinergic signal impacts bone through aging is also an important line of study that needs to be followed.

In males, increased mechanosensitivity clustered in the mid-physiological strain range, whereas females exhibited reduced mechanosensitivity at lower strain magnitudes. Factors determining osteocyte mechanosensitivity *in vivo* remain an active area of research. Key molecular players identified predominantly from *in vitro* studies include ion channels, integrins, connexins, and extracellular matrix organization(*66*–*69*). The prevailing hypothesis is that these elements collectively establish individual cellular thresholds for mechanosensitivity(*70*). Thus, it is plausible that deletion of Chrna1 or Rapsn alters the balance of osteocyte mechanosensitivity and/or modifies the strain threshold required to trigger cellular responses. Future research should explore the precise influence of osteocyte cholinergic receptor function on specific mechanotransduction pathways to clarify these relationships. An important implication of our findings is that modulation of cholinergic signaling can finely tune osteocyte mechanosensitivity, presenting a potentially powerful strategy for developing targeted clinical therapies for bone metabolic diseases.

Our anabolic mechanical loading studies revealed that cKO-driven deficiencies in female bone morphology could be rescued with tibial loading. Counterintuitively, *Rapsn*^Δ*Ocy/*Δ*Ocy*^ females also had suppressed bone formation and mineral apposition rate, indicating an alternative recovery mechanism. Since the present study only measured osteoblast-derived indices, we cannot disregard the possibility that alterations in osteoclast bone resorption are involved. ACh stimulation of MLO-Y4 cells increases the RANKL/OPG expression ratio 60-fold(*56*). Adding either mecamylamine, a non-competitive antagonist that inhibits all known nAChRs, or d-Tubocurarine, a nondepolarizing competitive antagonist that acts particularly on muscle nAChRs, ablates the response. Pyridostigmine, an AChE inhibitor that is not blood-brain barrier permeable (i.e. solely peripheral acting), increases RANKL/OPG serum levels and reduces femoral bone volume(*71*). Thus, nAChR activation on osteocytes may produce signals for osteoclast activity, implying *Rapsn* cKO could inhibit bone resorption. In contrast to females, any reductions in bone mass in the *Chrna1*^Δ*Ocy/*Δ*Ocy*^ males remained significantly lower even with loading, indicating distinct Ach-mediated bone metabolism regulation between the sexes.

Our 3-point bending mechanical tests indicated that *Chrna1* deletion in male osteocytes produced more ductile bone, however material analysis of the bone matrix did not indicate any changes to overall composition. These data suggest that any changes in mechanical behavior seen in male femurs likely comes from tissue arrangement. Further, they suggest an impact of osteocytes on tissue arrangement requires appropriate ACh signaling. Our studies here are limited in that we have used a constitutive model, which means any effect will be present through growth and development of bone tissue. To understand this phenomenon better, it would be helpful to use a temporally controlled cKO model in the future to allow bone tissue to develop normally.

Sex differences from cKOs imply different roles for ACh signaling, corroborating evidence that cholinergic signaling has a greater impact in females than in males. The negative bone measures associated with Alzheimer’s, which in general reduces ACh levels in the body, are mostly found in women and not men(*31*– *33, 71*). In general, women are more susceptible to both bone loss (i.e., post-menopausal osteoporosis) and dementia. Given the increased risk of osteoporosis with estrogen depletion, it may be possible that estrogen is also playing a role in the cholinergic regulation of bone, as indicated by the results in this study, although further investigation is necessary. Our results corroborate other studies on nAChRs in bone that found sex differences. For example, α7 global knockout does not produce changes in males(*72*), but in females, results in increased cortical area and thickness and reduced calcium content(*73*) as well as reductions in osteoclast count, RANKL serum levels, and increases to OPG serum levels(*74*). α9 global knockout in females leads to an increased number of apoptotic osteocytes(*75*). α2 global knockouts in females suffer from reduced trabecular bone volume and increased osteoclast count as well as resorption activity(*36*). Additional studies are needed to interrogate the potential for these additional subunits as osteocyte mechanotransduction regulators, and as sex-specific regulators of bone mechanotransduction more broadly.

We expect the osteocyte cholinergic receptor to have a similar conformation as muscle nAChRs, based on the component gene expression(*38*). These structures are heteropentameric with the stoichiometry (α1)_2_β1γδ (75). The α1-ε and α1-δ interfaces act as the binding sites for ACh, and the deletion of key hydrophobic regions to α1 abolish receptor function; thus, α1 knockout in osteocytes presumably inhibits nAChR function, although there may be stoichiometric substitution of the α subunit that we are not aware of that provide altered/limited receptor function. Rapsyn deletion permits healthy individual receptor formation while reducing overall signaling potency on account of increased receptor mobility. Rapsyn is also known to assist in faster recovery rates(*76*), meaning deletion may result in decreased signaling efficiency and effect. Future studies should include inducible deletion and pharmacological blockage of cholinergic signaling in bone to prevent any potential subunit substitutions. Follow-up RNA analyses would be helpful to identify any substitutions that occur with constitutive deletion.

*Chrna1* and *Rapsn* cKOs are unlikely to ablate cholinergic signaling entirely in the osteocyte, as other cholinergic receptors may be present(*38, 56*). Despite this, we have seen definitive evidence that osteocyte cholinergic receptors are important for bone development and response to mechanical challenges. This study did not trace the cholinergic source. This is important because ACh may be derived from a neuronal (e.g. parasympathetic/sympathetic nervous system(*36, 37*)) or non-neuronal (e.g. osteoblasts/bone lining cells(*37, 39*)) source, which has various biological implications and may also influence future therapeutic strategies. An important advance demonstrated in this study was the use of targeted genetic modification of ACh signaling in osteocytes specifically, rather than systemic pharmacological disruption or global knockout. Thus, we can show that ACh signaling to osteocytes directly impacts bone morphology and mechanoadaptation. This indicates an important new signaling axis, potentially implicating autonomic nervous system modulation of osteocyte activity.

With this work, we introduce a novel signaling pathway that suggests ACh can act on osteocytes *in vivo* to alter bone mass. Given the escalating prevalence of osteoporosis, investigating osteocyte-based therapeutics is a critical next step to advancing treatments beyond what is currently possible. Cholinergic neuromodulators such as AChE inhibitors are already commonly used to treat neurodegenerative diseases such as Alzheimer’s and have been gaining traction as potential therapeutics for bone diseases such as osteoporosis. To get closer to clinical translation, there is still a much-needed understanding of how cholinergic signaling impacts bone (specifically in osteocytes), and how this effect alters bone mechanotransduction. Our research is helping to elucidate the mechanism of cholinergic signaling in osteocyte mechanobiology and suggests a new signaling axis between brain and bone with far-reaching implications in bone biology that may be vital to treating bone disease.

## Materials and Methods

### All methods have been approved by Cornell Institutional Animal Use and Care Committee

#### Whole-Mount Tissue Clearing and Immunofluorescent Staining

We used a modified BABB (Benzoic Acid Benzyl Benzoate) tissue clearing approach. Mice were perfused with zinc formalin (contains 3.7% formaldehyde) and bones were dissected and fixed overnight in zinc formalin. Decalcification occurred over 5 days in 7.5 pH buffered 20% EDTA at 37 °C with daily solution changes and decolorized with 15% hydrogen peroxide for less than 1 hour. Immunofluorescent staining occurred before further proceeding. Samples are blocked overnight with 10% Donkey Serum (Sigma D9663-10ML) and 3% BSA in PBST. Primary antibodies are incubated at 1:100 dilution in a blocking buffer for 48 hours. Primary antibodies used were Rabbit Anti-Tyrosine Hydroxylase Antibody (TH, Sigma AB152) and Goat Anti-Vesicular Acetylcholine Transporter Antibody (VAChT, Sigma ABN100). Samples were washed in PBST overnight. Secondary antibodies were then incubated at 1:200 dilution in PBST for 48 hours. Secondary antibodies used were Alexa Flour 488 donkey anti-goat antibody (Thermo Fisher Scientific A11055) and Alexa Flour 546 donkey antirabbit antibody (Thermo Fisher Scientific A10040). Samples went through a final wash in PBST overnight. Tissue clearing was then finalized with dehydration in 2-ethoxyethanol for 4 hours, followed by 3 changes of acetonitrile between 4 hours or overnight at room temperature. Bones were then cleared in benzyl alcohol for 4 hours before being placed in a final refractive index matching solution of benzyl benzoate (refractive index 1.5) for long-term storage and imaging.

#### Light-Sheet Fluorescent Microscopy

Imaging of tissue-cleared third metatarsals (MT3s) was done using a Light-Sheet microscope (LaVision BioTec) at the Cornell Institute for Biotechnology Resource Center (Cornell BRC). Bones were mounted to clear hematocrit tubes using cyanoacrylate, then the tubes were attached to a custom-made mount for the microscope. Samples were completely submerged in a container filled with refractive index matching solution (Benzyl Benzoate and any residual Benzyl Alcohol; refractive index 1.5). Imaging was performed with two channels at excitation wavelengths of 488nm (6-8% power, emission collected with bandpass filter of 525/50) and 561 nm (9-11%, emission collected with bandpass filter 620/60) using an Olympus 2x CDC objective with a 3.5 collar and DBE correction for high RI. Two light sheets illuminated the sample in series from either the right or left directions. Images were acquired with a dynamic horizontal focus processing that blended the two images into one image, optimally reducing the scattering compared to a single light sheet. Both light sheets were calibrated and aligned in the z-plane before imaging. To image an entire bone, mosaic tiling with a 10% overlap and 1×3 tiles were used with 3.2x zoom, sheet NA 1.20, and sheet width 40%. The final pixel resolution was 0.94×0.94×4μm. Files were saved in TIF OME format for each tile and channel, comprising 6 raw data images. Images are stitched using BigStitcher in FIJI (NIH). Stitched images were converted into Imaris (Oxford Instruments) files. Channel shifting, 100μm radius background subtraction, and 1.2 gamma correction are applied in Imaris. Finally, using the filaments tool in Imaris, large nerve fiber-like objects were manually traced and quantified using Imaris metrics.

#### *In vivo* Loading of the Third Metatarsal

Our group uses an innovative approach to observe live osteocyte cell and molecular level fluorescent signals *in vivo*. Additionally, we can collect these observations with simultaneous tissue level mechanical loading (**Fig S6**). This approach is an extremely powerful approach for studying basic mechanobiology *in vivo*. Before loading, calibrations were conducted on cadaveric mouse MT3 with digital image correlation (DIC) to determine position-controlled loading regimes with corresponding strain values. Hind paws from 3-6 mice for each mouse group were amputated and stored in saline-soaked gauze at −20°C until calibration. For calibration, paws were thawed at room temperature in DPBS. The MT3 was surgically isolated as described elsewhere and a 1mm diameter cylindrical fulcrum pin was inserted just underneath, separating the bone from any muscle fascia and soft tissue(*30*). A bracket for the two side pins was then positioned over the paw and connected with a screw to the load cell and actuator. Calibration was performed for each genotype according to previously published work(*30*).

For experiments, mice were initially placed under anesthesia in a small container with a 1 L/min flow rate and 3% isoflurane. Mice were then transferred onto the loading device stage with a heating pad and head covering for more direct anesthesia, where the isoflurane was maintained at 1.5-2%, at which point surgery in the same manner as with calibration took place on the left hind paw. With a live mouse, it was important to avoid the rupture of blood vessels and to remove any tendon that resides longitudinally across the metatarsals. The fulcrum pin and bracket were attached accordingly. The paw was placed in a room-temperature DPBS bath for the duration of the experiment.

#### Nerve-Osteocyte Proximity Analysis *In vivo*

A total of six ChAT strain mice (4 males and 2 females) were used in the study, all 10 months old. For labeling the osteocytes in the MT3 bone, RGD-functionalized integrin targeting nanoparticles (RGD C-Dots) (15µL of 10µM) were injected via subcutaneous method in the right and left paws, respectively. Following injection, 5-minute incubation was maintained for RGD C-Dots. Imaging was performed under anesthesia using a two-photon microscope (Thor Labs Bergamo II; Newton, NJ) and a 20x water immersion objective with a long working distance (XLUMPLFLN, Olympus) to assess fluorescence signals, with specific observations on ChAT expression and RGD C-Dots localization. The RGD C-Dots fluorescence was examined at multiple wavelengths, with stronger signals detected at 1020nm.

Images containing nerve fibers and osteocytes were processed by first confirming file paths and loading corresponding binary masks. Osteocytes were identified by converting images to the HSV color space and applying a dual-range threshold to isolate red-stained osteocyte structures, followed by contour extraction and centroid calculation. A secondary mask was applied to remove image regions dominated by background noise. Osteocytes overlapping nerve fibers or these masked noisy regions were excluded to enhance quantification accuracy. Additionally, osteocytes located within 10 µm of each other were filtered to prevent duplicate counting. Nerve fiber boundaries were extracted from binary masks, and the shortest Euclidean distance from each osteocyte centroid to the nearest nerve fiber boundary was calculated using OpenCV’s cv2.pointPolygonTest. Pixel distances were converted to microns based on image scale bars. Each osteocyte was assigned to its nearest nerve fiber, and data including osteocyte coordinates, nerve assignments, and distances were recorded. Finally, osteocytes were visually labeled and color-coded according to their nearest nerve fiber for validation purposes.

Given the large osteocyte sample size (n > 200), the Kolmogorov-Smirnov test was employed to assess normality of data distributions separately for males and females(*77*). Non-parametric distributions (p < 0.05) were identified in both groups; therefore, statistical comparisons between males and females were conducted using the Mann-Whitney U test. Statistical significance was defined as p < 0.05.

#### Intravital Imaging of Calcium Dynamics with Two-Photon Microscopy

*In vivo* fluorescence signals of GCaMP6f from Ca^2+^ dynamics in osteocytes were visualized with a two-photon microscope (Thor Labs Bergamo II; Newton, NJ) and a 20x water immersion objective with a long working distance (XLUMPLFLN, Olympus). Osteocytes were imaged at the dorsal mid-shaft region of the MT3, directly above where the fulcrum pin was placed beneath the bone, and beneath the periosteal surface, verified with visible autofluorescence from the bone surface. Cyclic haversine position-controlled loading at 1 Hz was applied for 60 seconds to previously calibrated peak strains of 250, 500, 1000, 2000, and 3000με. These levels encompass the range of strains that have been reported during physiological activities from *in vivo* strain gage studies, with strains up to 2000με characteristic of habitual activities, while strains on the order of 3000με are seen in extreme activities(*4, 78*). A 15-minute wait time to account for osteocytes’ refractory period was instigated between each imaging/loading session(*79*). The ROI was 512 × 512 pixels with 2x zoom, leading to a pixel size of 0.545 × 0.54μm. Excitation was at 920 nm wavelength from a tunable Ti-Saphire laser (Coherent Chameleon Discovery NX) with a power selected such that the final output from the objective was between 30-35mW. A photomultiplier (PMT) with a 490-560nm bandpass filter was used for detection with a voltage gain of 0.1-12V. Time series images were acquired by averaging 3 frames with a sampling rate of 29.4Hz, leading to a final frame rate of 9.8fps. Across an approximate 150-second period, 1500 8-bit TIF images were collected, with 60 seconds of non-loading, 60 seconds of loading, and 30 seconds of resting.

#### MT3 Loading Image Analysis

Intensity measurements were performed by postprocessing images using ImageJ. In-plane movements in the x and y were removed using a template-matching plugin. ROIs were selected with a semi-automated process. Gaussian blur with a sigma of 1 was applied to smooth edges. Automated size exclusion was performed, filtering ROIs for only 150-650 pixel sizes. Finally, users would manually make final adjustments to ROIs by eye, deleting and adding ROIs as needed that were not properly captured by thresholding. An ROI indicating the background autofluorescence was included in an area where no cells were present. The average pixel intensity for each ROI was collected across all 1500 time series frames and output as a CSV.

In MATLAB, ROIs that exceeded a cutoff of the mean background intensity plus 3 standard deviations were included in the final analysis as fluorescent osteocytes. The intensities for each cell of interest were detrended linearly to correct for photobleaching and normalized to the mean intensity for that cell over 40 seconds before loading. Peaks and troughs were identified using islocalmax and islocalmin in MATLAB with a prominence of 0.01. Amplitudes across the non-loaded and loaded intervals were computed by subtracting coupled troughs from peaks. Changes in amplitude fluorescent intensity were computed as follows: ΔF/F=(F_load-F_nonload)/F_nonload. F_load indicates the amplitude during loading and F_nonload indicates the amplitude without loading. Responding cells were defined as cells showing a > 25% increase in ΔF/F. ΔF/F was then plotted for responding cells only. Of 9222 ROIs (indicating cells with the possibility of repeats across different loading strains), 25 ROIs were removed as outliers with a change in fluorescent intensity greater than what we determine here as the dynamic range (>13) of GCaMP6f with repeated stimulation(*80*).

#### Generation of *Chrna1* and *Rapsn* cKO mouse lines

*Chrna1* and *Rapsn* deletion from the mouse genome was conducted by purchasing tm1a-targeted embryonic stem cells for both *Chrna1* and *Rapsn*, from EUCOMM, which were expanded and injected into blastocysts, returned to pseudo-pregnant females, and screened for germline transmission in pups. Both *Chrna1* and *Rapsn* germline deletion were lethal, consistent with previous studies(*55, 81*), so we took steps to generate Cre-ready conditional alleles (tm1c) for both loci (**Fig S1**). Briefly, the correctly targeted mice harbor LacZ and Neo cassettes in intron 2 (for *Rapsn*) or 3 (for *Chrna1*) flanked by flippase recognition target (FRT) sequences. The Neo cassette was flanked by loxP, and a third loxP was introduced after the final exon in the targeted sequence (exon 4 for *Chrna1* and exon 5 for *Rapsn*). To convert the tm1a allele to a tm1c conditional floxed allele, tm1a mice were crossed to Rosa26Flp mice (Jax stock 012930; described elsewhere(*82*)) to induce germline recombination of the FRT sites and delete the LacZ/Neo/5’loxP insert, leaving one lonely FRT site and the targeted exon(s) flanked by loxP. These tm1c mice were then crossed to the Dentin matrix protein 1 Cre (10kb Dmp1-Cre; Jax stock 023047) to induce conditional deletion of *Chrna1* exon 4 or *Rapsn* exons 3-5 (both target regions house crucial exons) in osteocytes and late-stage osteoblasts. The Dmp1-Cre line has been described elsewhere(*83*). *Chrna1* and *Rapsn* lines were validated for correct targeting by long-range PCR using primer pairs spanning the homology arms at both 5’ and 3’ ends of the targeting region. Both mutant mouse lines were on a pure C57BL/6J background. Female and male mice with Dmp1-Cre-positive conditional knockouts (cKO) in osteocytes are labeled *Chrna1* cKO and *Rapsn* cKO, with littermate Dmp1-Cre-negative controls *Chrna1*^+*/*+^ and *Rapsn* ^+*/*+^ used as the respective reference.

#### Generation of osteocyte-targeted GCaMP6f mice with *Chrna1* and *Rapsn* cKO

GCaMP6f is a recombinant protein construct containing calmodulin (CaM), the CaM binding myosin light chain kinase fragment (M13), and enhanced green fluorescent protein (EGFP). Bright EGFP fluorescence is observed following Ca^2+^ binding to CaM, causing a subsequent binding of CaM to M13 and a conformational change that results in increased fluorescence intensity. Mice exhibiting osteocyte-targeted expression of GCaMP6f were achieved by crossing Ai38 mice B6J.CgGt(ROSA)26Sortm95.1(CAGGCaMP6f)Hze/MwarJ (JAX Labs, strain no. 028865), which contain GCaMP6f DNA behind a Lox-STOP-Lox codon(*80*), with DMP1-Cre mice [B6N.FVB-Tg(Dmp1-cre)1Jqfe/ BwdJ; (JAX Labs, strain no. 023047) (*83*). Both parent strains are on a C57BL/6 background. Previous studies with this mouse model used GCaMP3f and validated fluorescent expression in cortical osteocytes(*30, 84*). Our substitution with GCaMP6f was motivated by the fact that GCaMP6f was created as an ultrasensitive protein calcium sensor that outperforms previous GCaMP iterations with a variant that has faster kinetics and greater dynamic range(*80*).

Finally, to obtain homozygous knock-in of GCaMP6f with simultaneous knock-out of either *Chrna1* and *Rapsn*, we crossbred GCaMP6f mice with each conditional knockout three generations deep to obtain GCaMP6f-*Chrna1* cKO and GCaMP6f-*Rapsn* cKO mice. In this study, intact GCaMP6f mice were used as controls. For experiments, male and female mice were aged until skeletal maturity (16-18 weeks).

#### *In vivo* Tibial Loading

The axial tibial compression model was applied to Cre-positive and Cre-negative mice as previously described(*85, 86*). Before loading, calibration mice for each sex and genotype were sacrificed at 17 weeks of age to collect strain measurements. Disarticulated hindlimbs previously stored at 20°C were used for strain gauge calibration. A single-element strain gauge (EA-06-015DJ-120; Vishay Precision Group, Malvern, PA, USA) was applied to the posterior midshaft surface of the tibia. The load-strain relationship was measured for each sample by applying step loading up to 11 N. A load-strain curve was derived from the simultaneous recording of the voltage output from the load cell and strain gauge. All tests were averaged within each genotype/sex to establish a calibration curve.

For *in vivo* anabolic loading, mice were anesthetized using isoflurane inhalation, and their right hindlimb (knee to paw) was loaded to a calibrated peak strain value of 2250 µε using a haversine waveform (2 Hz, 180 cycles). Mice were given three bouts over 5 days with a day of rest between each bout. Mice were sacrificed 12 days after the final bout.

#### Dual-Energy X-Ray Absorptiometry (DEXA)

Bone mineral content (BMC) of the 3^rd^-5^th^ lumbar vertebrae, the right leg, and the whole body was evaluated *in vivo* using dual-energy x-ray absorptiometry (DEXA). Mice were anesthetized via inhalation of 2.5% isoflurane (IsoFlo; Abbott Laboratories, North Chicago, IL) mixed with O_2_ (1.5 liter/min) for approximately 8 minutes, including induction and scanning. The mice were placed prone on a specimen tray within the scanner. Bone mineral density (BMD) was computed using raw bone mineral content for each region of interest. Scans were performed at 4, 6, 9, 13, 17, and 21 weeks of age. All BMC measures were normalized by body weight to eliminate the confounding effects of changing body size and weight during growth.

#### Microcomputed Tomography (µCT)

Tibiae were harvested 12 days after the final bout of tibial loading, wrapped in saline-soaked gauze, and stored in an airtight container at −20°C. Geometric properties of trabecular and cortical bone were evaluated using high-resolution microcomputed tomography (µCT, Scanco Medical AG). Distal and mid-diaphysis 9 µm voxel size resolution scans were imported into Scion Image version 4.0.2 (Scion Corporation), where properties were calculated using standard and customized macros. Standard measurements included cortical area (mm^2^), maximum (Imax, mm^4^) and minimum (Imin, mm^4^) cross-sectional moments of inertia, polar moment of inertia (pMOI, mm^4^), total volume (TV, mm^3^), bone volume (BV, mm^3^), bone volume fraction (BV/TV, %), connectivity density (Conn.D, mm^*−*3^), structure model index (SMI), trabecular number (Tb. N, mm^*−*1^), trabecular thickness (Tb. Th, µm), and trabecular separation (Tb. Sp, µm), bone mineral content (BMC, g), and bone mineral density (BMD, mg/cm^3^).

#### Bone Histomorphometry

An intraperitoneal injection of calcein was given 1 day after the final bout of tibial loading, followed by an intraperitoneal injection of alizarin complexone 8 days later. Mice were sacrificed 3 days after alizarin injection. The right and left tibias were harvested and placed in 10% neutral buffered formalin for 2 days, then stored in 70% ethanol at 4 °C. Tibiae and femora were dehydrated in graded alcohols, cleared in xylene, and embedded in methyl methacrylate. To measure load-induced bone formation, thick sections were cut from the tibia approximately 3 mm proximal to the tibiofibular junction and ground down with a polisher to approximately 50µm (Multiprep Polishing System Allied High Tech Products). Tibial diaphysis sections were mounted unstained to visualize and read the calcein and alizarin labels administered at 17 and 18 weeks. Sections were imaged on a fluorescent microscope using filter sets that provide excitation and emission for the calcein and alizarin wavelengths. Digital images were imported into ImageJ (1.54f NIH, Java 1.8.0_322) and the following histomorphometric measurements(*87, 88*) were recorded for the periosteal and endosteal surfaces: total perimeter, single-label perimeter (sL.Pm), double-label perimeter (dL.Pm), interlabel thickness (Ir.L.Th), total bone area, and marrow area. The following results were calculated: mineral apposition rate (MAR = Ir.L.Th/8 days), mineralizing surface (MS/BS = (0.5 × sL.Pm + dL.Pm) / total perimeter × 100), and bone formation rate (BFR/BS = MAR × MS/BS × 3.65). Relative formation parameters for loading effects were calculated for each mouse by subtracting the non-loaded (left tibia) values from the loaded (right tibia) values.

#### Mechanical Testing

Mice were sacrificed at 16-18 weeks and the right femurs were harvested and stored at −20°C in saline-soaked gauze until testing. On the day of testing, femurs were brought to room temperature in a saline bath for over 1 hour. Femurs were positioned posterior side down across the two lower supports of a three-point bending fixture. The femurs were loaded to failure in monotonic compression with a span length of 9 mm using a crosshead force and displacement measurements were collected at every speed of 0.2 mm/s using a mechanical testing system (Bionix; MTS, Eden Prairie, MN, USA). Force and displacement measurements were measured using a 25-pound load cell (SSM-25 Transducer Techniques, Temecula, CA, USA) and a linear variable differential transducer (a calibration curve spanning the expected force range was generated before testing). From the force versus displacement curves, ultimate force (N), ultimate energy (mJ), failure energy (mJ), stiffness (N/mm), yield force (N), and Post-Yield Displacement (PYD, mm) were calculated using standard equations(*89*). Before testing, the length of each femur was measured to the nearest 0.01 mm, along the diaphyseal axis using digital calipers.

#### Fourier Transform Infrared Spectroscopy (FTIR)

Fourier Transform Infrared Spectroscopy (FTIR) was used to evaluate bone compositional properties. Left femora from mice used for mechanical testing were harvested and stored at −20°C in saline-soaked gauze until specimen preparation(*90*). Briefly, epiphyses of the femur were detached and bone marrow was removed using a centrifuge. The pieces were defatted in 3×15 minute solution changes of 100 % isopropyl ether. Samples were lyophilized in a vacuum drier and made into powder using a CryoMill (770; SPEX SamplePrep, Metuchen, NJ, USA). Pellets were constructed by mixing 2.0-3.0 mg of powdered bone with dry potassium bromide to make a 200 mg mixture. Pellets were pressed with a 13-mm-diameter die and a maximum load of 10 tons using an evacuable pellet press (Pike Technologies, Fitchburg, WI, USA). FTIR spectra were collected at a spectral resolution of 4 cm^-1^ over the spectral range of 800 to 2000 cm^-1^ using an FTIR spectrometer (Spotlight 400; Perkin-Elmer Instruments, Waltham, MA, USA). Spectra were analyzed in which areas were defined between the following peaks for Phosphate: 916-1180 cm^*−*1^, Amide I: 1596-1712 cm^*−*1^, and Carbonate: 852-890 cm^*−*1^. The peak height ratios were defined as Crystallinity: 1030/1020, Acid Phosphate: 1126/1096, and collagen maturity (cross-linking ratio): 1660/1690.

#### Statistical Analysis

The effect of the cholinergic cKO on measurements of bone geometry, mineral density, mechanical properties, and metabolism was determined using a One-way ANOVA with group as the factor (Cre-negative littermate controls against Cre-positive experimental mice for the same sex) with a Tukey’s post-hoc analysis followed by unpaired one-tailed T-tests with alternative hypotheses as less than or greater than the control group depending on our hypothesis regarding the changing phenotypes. For DEXA BMD measurements, Two-way ANOVA was performed with interactions between Genotype and Age. A Sidak post-hoc test was computed across genotypes for each time point. Outliers greater than three standard deviations from the mean for each metric grouped by sex were removed. Statistical tests were conducted using R or Prism with α = 0.05.

The effect of the cholinergic cKO on the percent cells responding and change in fluorescence intensity was determined using a two-way ANOVA with a group (GCaMP controls vs GCaMP cKOs) and strain (250, 500, 1000, 2000, 3000με) as factors followed by unpaired one-tailed T-tests with alternative hypotheses as less than or greater than the control group depending on our hypothesis regarding the changing phenotypes. Statistical tests were conducted using R or Prism with α = 0.05.

The mixed effects of sex and genotype within the dynamic histomorphometry, mechanical testing, microCT, and *in vivo* loading datasets was assessed using a mixed model run in RStudio version 2023.12.1+402 using the lme4 package. Resulting tables are in the Supplementary Materials.

Sample size varied by group and assay. The sample sizes for each group (CHRNA:C, RASPN:R) are as follows: DEXA (C: 6-13, R: 3-7), Dynamic Histomorphometry (C: 8-14, R: 7-12), FTIR (C: 3-6, R: 5-7), Mechanical Testing (C: 9-14, R: 9-12), MicroCT (C: 9-22, R: 4-8), Skeletal Innervation (C: 3, R: 3-4), MT3 Loading (GCaMP: 7-9, C: 9-10, R: 12-14).

## Acknowledgments

We thank Karly Hooper for maintaining mouse colonies for use in these experiments, along with the entire CARE staff at Weill Hall. We thank the Donnelly Lab in the Materials Science and Engineering Department at Cornell University for assistance with FTIR collection. We thank the Schaffer-Nishimura lab for providing ChAT mice. We thank Damien Laudier for histological guidance for the bone-clearing protocol reported here. We thank the Wiesner Lab in the Material Science and Engineering Department for providing C’Dots for osteocyte imaging.

## Funding

National Science Foundation 2418809 (KJL)

NIH 5U24AT011969-03 (Pilot Award from a parent U24) (KJL)

NIH S10OD023466 (N/A)

Mong Fellowship, Cornell NeuroTech (MMA)

Ford Foundation Fellowship, National Academy of Science (MMA)

## Author contributions

Conceptualization: KJL MMA

Methodology: KJL MMA AS MO MDM

Data Curation: KJL MMA AS AGO

Formal Analysis: AS MMA AGO

Software: AS MMA

Investigation: MMA KJL MDM MW AS AGO MO AGO

Visualization: MMA KJL AGO MDM

Validation: MMA AGO

Supervision: KJL

Writing—original draft: KJL MMA AGO

Writing—review & editing: KJL AGO MDM

## Competing interests

KJL and MDM are inventors on a patent related to this work filled by Cornell University (PCT/US24/15289 filed on February 9, 2024, Published August 15, 2024). The authors declare no other competing interests.

## Data and materials availability

All data are available in the main text or the supplementary materials.

## Notes

### Competing Interest Statement

The authors have declared no competing interest.

### Summary of Updates

We have made edits to figures, text, and added additional data to strengthen the manuscript's impact.

