## Supplemental Figures for "Osteocyte Nicotinic Acetylcholine Receptors Impact Bone Mechanoadaptation in a Sexually Dimorphic Manner"

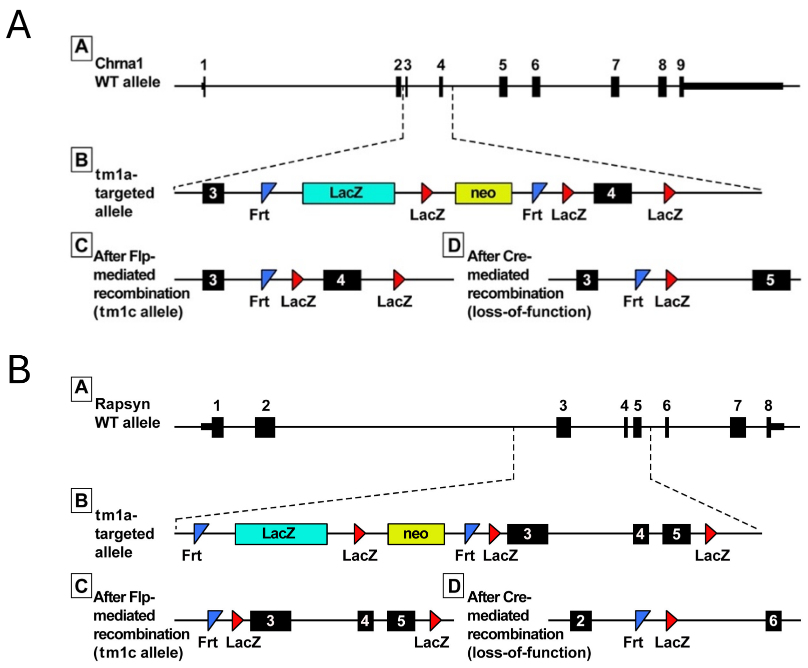


**Supplemental Figure 1**: Cre-Lox based genetic modification strategy. Schematic showing the floxed (A) *Chrna1,* (B) *Rapsn* loci (2 exons), and the location within exon 1 of the forward and reverse primers used to probe for the intact floxed allele in genomic DNA from cortical bone.

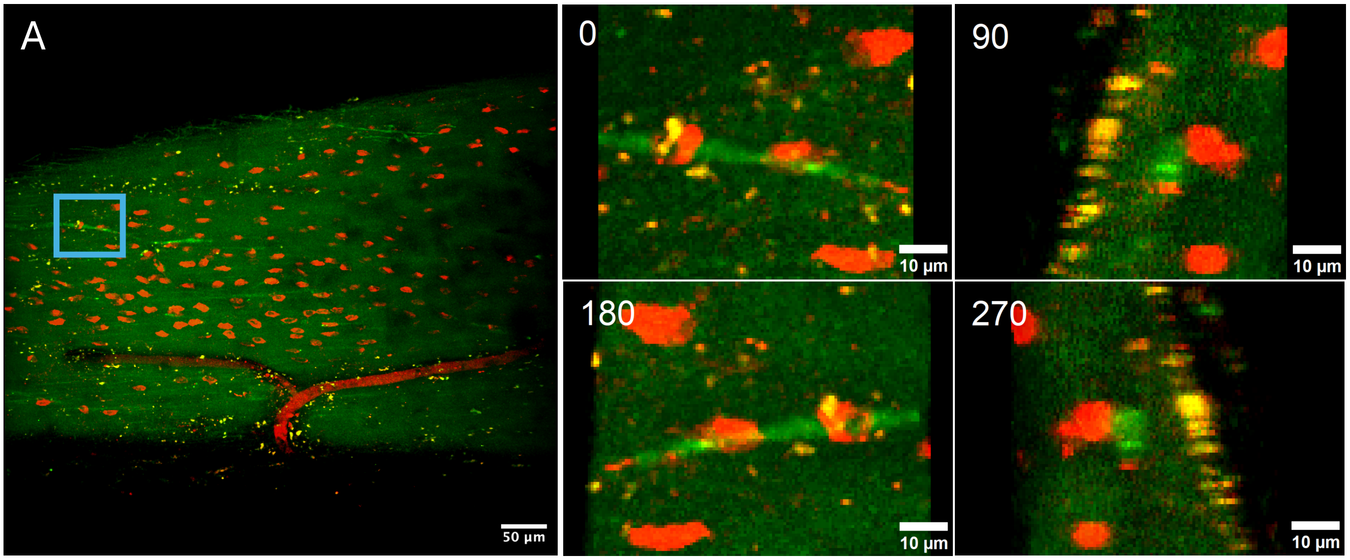


**Supplemental Figure 2**: (A) Intravital Z-stack image (40μm) of metatarsal bone osteocytes in mice expressing GCaMP6f in acetylcholine transferase expressing cells taken using two photon microscopy at about 20μm below the periosteal surface. Red fluorescence signal is driven by a 15μL volume of ultrasmall RGD-functionalized integrin targeting nanoparticles (5-7nm) at 10μM concentration. The nanoparticles were injected subcutaneously at the volar aspect of the hind paw and incubated for 45 minutes. After the incubation period, the third metatarsal was surgically exposed and imaged within 5 additional minutes. This static 3D image indicates colocalization of osteocytes, labeled in red, with cholinergic nerve fibers, labeled in green. Images taken at 90-degree rotation intervals in an ROI are included, indicating a direct interaction between osteocytes and nerve fibers.

**Normal distribution Test: Kolmogorov-Smirnov Test**

| | Male | Female |
|---|---|---|
| KS distance | 0.0803 | 0.1011 |
| P value | <0.0001 | <0.0001 |
| Passed normality test (α=0.05)? | No | No |
| P value summary | **** | **** |

**Male vs Female: Mann Whitney Test**

| | |
|---|---|
| P value | 0.0011 |
| P value summary | ** |
| Significantly different (P < 0.05)? | Yes |
| One- or two-tailed P value? | Two-tailed |

**Supplemental Figure 3**: Statistical tests for examining osteocyte distributions within 100µm of cholinergic nerves *in vivo*. Kolmogorov-Smirnov test was used for normality assessment. Mann Whitney non-parametric test was used for comparing between sexes.

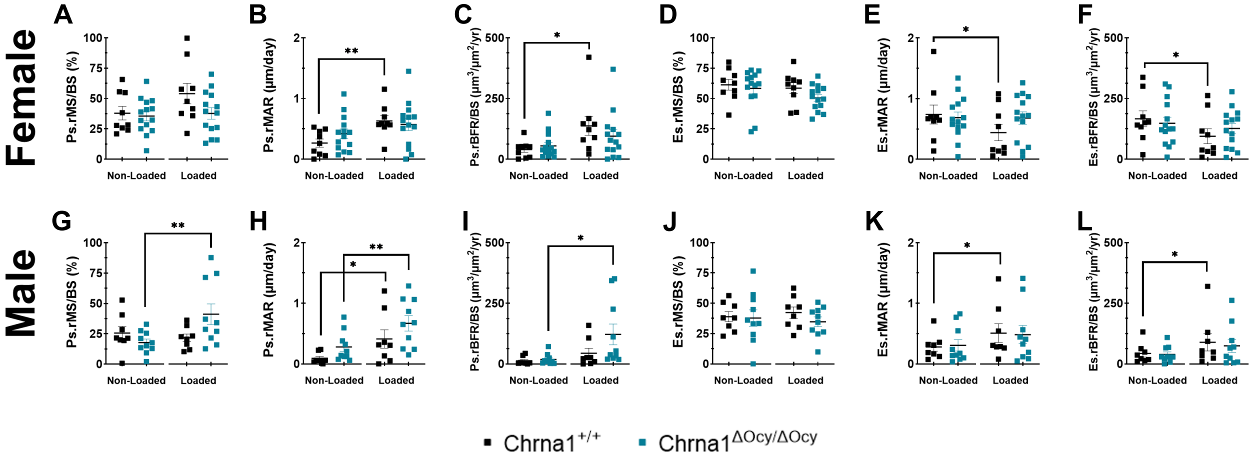


**Supplemental Figure 4**: *Chrna1* cKO impaired the anabolic response to mechanical loading. Fluorochrome labeling was used for osteoblast-derived indices of dynamic histomorphometry. Loaded and non-loaded contralateral limbs are compared here. Indices were computed for both the endosteal and periosteal surfaces. Generally, loaded control limbs showed anabolic bone growth for both males and females. Cre-negative *Chrna1* controls showed differences between the non-loaded and loaded limbs for Ps.rMAR, Es.rMAR, and Es.rBFR/BS in males and females (B, E-F). There were no differences between the non-loaded and loaded limbs in the *Chrna1* cKO females (A-F). In male *Chrna1* cKO, Ps.rMS/BS, Ps.rMAR, and Ps.rBFR/BS were increased between non-loaded and loaded limbs (G-I). Unpaired Student T-tests were performed between the non-loaded and loaded limb within sex and genotype (*$p<0.05$). N= 8-22 per group and assay.

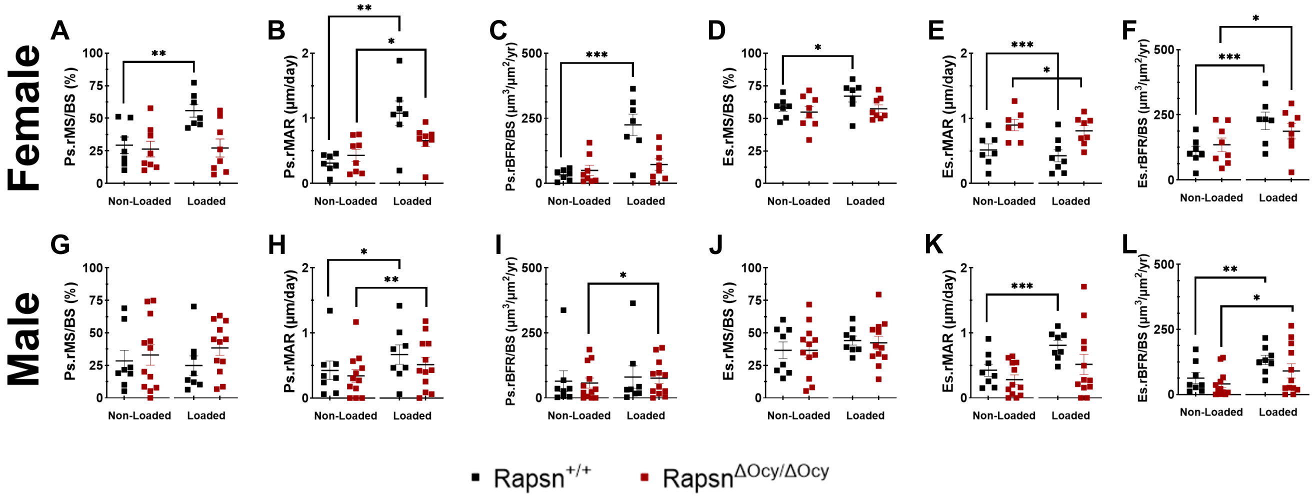


**Supplemental Figure 5**: *Rapsn* cKO impaired bone formation in response to mechanical load. Fluorochrome labeling was used for osteoblast-derived indices of dynamic histomorphometry. Indices were computed for both the endosteal and periosteal surfaces. Loaded and non-loaded contralateral limbs are compared here. Indices were calculated for both the endosteal and periosteal surfaces. Generally, loaded control limbs showed anabolic bone growth for both males and females. In female *Rapsn* cre-negative controls, the non-loaded and loaded limbs differed in all measurements (A-F). *Rapsn* cKO females non-loaded to loaded limbs varied for Ps.rMAR, Es.rMAR, and Es.rBFR.BS (B, E-F). Male *Rapsn* cre-negative controls showed increases between the non-loaded and loaded limbs for Ps.rMAR, Es.rMAR, and Es.rBFR/BS (H, K-L). The male *Rapsn* cKO increased for Ps.rMAR, Ps.rBFR/BS, and Es.rBFR/BS (H-I, L). Unpaired Student T-tests were performed between the non-loaded and loaded limb within sex and genotype (*p<0.05). N = 4-12 per group and assay.

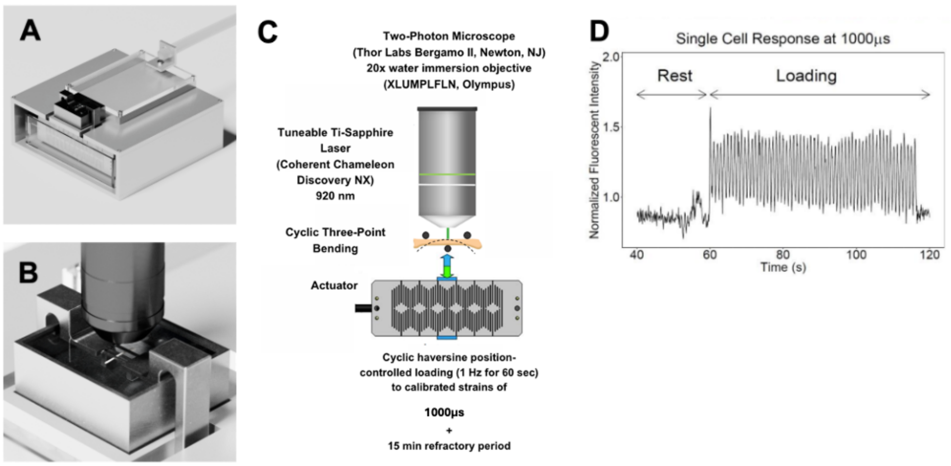


**Supplemental Figure 6**: Synopsis of metatarsal *in vivo* loading with intravital imaging. A) The custom-built device consists of a platform for anesthetized mice to rest on, a piezo actuator beneath a PBS water bath, and a loading bracket. (B) The device was designed to provide space for a microscopic objective to nest within the loading bracket for simultaneous microscopic observation of cells within bone tissue while loading occurs. (C) Three-point bending is inverted, with loading occurring at the proximal and distal ends of the third metatarsal. This conformation allows for the maximum strain and minimum displacement to be co-localized. It also exposes this site to the microscope objective. (D) Representative $Ca^{2+}$ signaling data from an osteocyte *in vivo*.

Supplemental Table 1. Full microCT dataset for female *Chrna1* cKO mice tibias. P-values <0.05 are bolded.

| Female CHRNA1 Tibia | | | | | | | | | | |
|---|---|---|---|---|---|---|---|---|---|---|
| | Non-loaded | | | | | Loaded | | | | |
| | Cre- | St Dev | Cre+ | St Dev | p Value | Cre- | St Dev | Cre+ | St Dev | p Value |
| **TrabTV** | 2.6748 | 0.2048 | 2.6202 | 0.1651 | 0.2023 | 2.7759 | 0.1444 | 2.6641 | 0.1004 | **0.0110** |
| **TrabBV** | 0.0411 | 0.0164 | 0.0565 | 0.0197 | **0.0104** | 0.0487 | 0.0144 | 0.0505 | 0.0207 | 0.3911 |
| **TrabBVTV** | 0.0127 | 0.0050 | 0.0188 | 0.0073 | **0.0044** | 0.0142 | 0.0039 | 0.0163 | 0.0073 | 0.1658 |
| **TrabConnD** | 4.8474 | 5.3775 | 6.7517 | 5.1949 | 0.1662 | 3.9066 | 2.8155 | 7.6355 | 7.8709 | 0.0517 |
| **TrabSMI** | 3.6034 | 0.3630 | 3.5654 | 0.3926 | 0.3858 | 3.6817 | 0.2893 | 3.6038 | 0.3276 | 0.2419 |
| **TrabTbN** | 2.2214 | 0.3623 | 2.4841 | 0.2643 | **0.0115** | 2.2782 | 0.3282 | 2.4491 | 0.2563 | 0.0501 |
| **TrabTbTh** | 0.0319 | 0.0034 | 0.0333 | 0.0027 | 0.1041 | 0.0346 | 0.0042 | 0.0331 | 0.0048 | 0.1752 |
| **TrabTbSp** | 0.4497 | 0.0903 | 0.4075 | 0.0457 | **0.0474** | 0.4249 | 0.0404 | 0.4107 | 0.0459 | 0.1823 |
| **TrabDenTV** | 41.0504 | 13.8765 | 51.6002 | 15.3074 | **0.0234** | 45.7923 | 17.2003 | 47.5769 | 14.1884 | 0.3741 |
| **TrabDenBV** | 839.5432 | 22.7841 | 857.1475 | 33.0945 | **0.0434** | 847.6324 | 21.3185 | 849.0279 | 25.9650 | 0.4359 |
| **TrabBMC** | 0.0346 | 0.0141 | 0.0487 | 0.0179 | **0.0088** | 0.0419 | 0.0140 | 0.0431 | 0.0183 | 0.4194 |
| **TotTV** | 5.6372 | 0.1707 | 5.6767 | 0.1421 | 0.2476 | 5.6982 | 0.2004 | 5.7332 | 0.2709 | 0.3357 |
| **TotBV** | 2.0827 | 0.2543 | 2.2341 | 0.3268 | 0.0707 | 2.0852 | 0.2653 | 2.2046 | 0.3224 | 0.1234 |
| **TotBVTV** | 0.3699 | 0.0479 | | | | 0.3658 | 0.0435 | 0.3833 | 0.0430 | 0.1240 |
| **TotDenTV** | 374.6287 | 84.9203 | 401.3746 | 79.3465 | 0.1749 | 367.5884 | 75.7051 | 396.7192 | 78.0270 | 0.1387 |
| **TotDenBV** | 1000.2799 | 99.6627 | 1021.8455 | 92.8794 | 0.2593 | 995.8831 | 93.1876 | 1023.6180 | 93.9947 | 0.1970 |
| **TotBMC** | 2.1054 | 0.4638 | 2.3101 | 0.5319 | 0.1203 | 2.0963 | 0.4502 | 2.2833 | 0.5254 | 0.1367 |
| **CortTV** | 5.6372 | 0.1707 | 5.6767 | 0.1421 | 0.2476 | 5.6982 | 0.2004 | 5.7332 | 0.2709 | 0.3357 |
| **CortBV** | 2.0252 | 0.2373 | 2.1776 | 0.3195 | 0.0621 | 2.0295 | 0.2512 | 2.1436 | 0.3043 | 0.1209 |
| **CortBVTV** | 0.3599 | 0.0463 | 0.3765 | 0.0451 | 0.1615 | 0.3561 | 0.0414 | 0.3727 | 0.0401 | 0.1215 |
| **CortBMC** | 2.0558 | 0.4468 | 2.2614 | 0.5253 | 0.1140 | 2.0471 | 0.4374 | 2.2287 | 0.5066 | 0.1358 |
| **CortBMD** | 1.0047 | 0.1030 | 1.0260 | 0.0949 | 0.2685 | 0.9992 | 0.0950 | 1.0277 | 0.0960 | 0.1952 |
| **MSMeanpMOI** | 0.3205 | 0.0541 | 0.2966 | 0.0460 | 0.0939 | 0.3044 | 0.0591 | 0.3073 | 0.0521 | 0.4405 |
| **MSMeanImax** | 0.2131 | 0.0386 | 0.1912 | 0.0311 | **0.0436** | 0.1996 | 0.0390 | 0.2025 | 0.0384 | 0.4164 |
| **MSMeanImin** | 0.1074 | 0.0177 | 0.1054 | 0.0157 | 0.3703 | 0.1047 | 0.0209 | 0.1024 | 0.0116 | 0.3538 |
| **MSMeanBArea** | 0.7632 | 0.1144 | 0.7189 | 0.1087 | 0.1283 | 0.7360 | 0.1469 | 0.7285 | 0.1027 | 0.4317 |
| **MSMeanTArea** | 1.6045 | 0.0628 | 1.6134 | 0.0939 | 0.3832 | 1.6241 | 0.0893 | 1.6166 | 0.0932 | 0.4116 |
| **MsTbTh** | 0.1822 | 0.0251 | 0.1728 | 0.0238 | 0.1333 | 0.1759 | 0.0340 | 0.1744 | 0.0207 | 0.4352 |
| **MsDenTV** | 519.5902 | 89.4316 | 489.9269 | 79.8422 | 0.1576 | 515.3836 | 92.5341 | 490.4632 | 74.6326 | 0.1969 |
| **MSDenBV** | 1136.1930 | 59.5752 | 1112.9043 | 57.5873 | 0.1275 | 1132.0129 | 68.1698 | 1111.0863 | 56.6134 | 0.1688 |
| **MSDenBVTV** | 2.2338 | 0.2930 | 2.3111 | 0.2609 | 0.2113 | 2.2447 | 0.2884 | 2.2994 | 0.2407 | 0.2763 |

Supplemental Table 2. Full microCT data for male *Chrna1* cKO mice tibias. P-values <0.05 are bolded.

**Male CHRNA1 Tibia**

| | Non-loaded | | | | | Loaded | | | | |
|---|---|---|---|---|---|---|---|---|---|---|
| | Cre- | St Dev | Cre+ | St Dev | p Value | Cre- | St Dev | Cre+ | St Dev | p Value |
| **TrabTV** | 3.4095 | 0.5253 | 3.4057 | 0.5655 | 0.4927 | 3.3878 | 0.5696 | 3.4502 | 0.6050 | 0.3890 |
| **TrabBV** | 0.2994 | 0.1344 | 0.3098 | 0.1730 | 0.4313 | 0.2946 | 0.1275 | 0.2931 | 0.1412 | 0.4884 |
| **TrabBVTV** | 0.0816 | 0.0314 | 0.0830 | 0.0381 | 0.4577 | 0.0815 | 0.0308 | 0.0783 | 0.0322 | 0.3935 |
| **TrabConnD** | 56.5814 | 31.6758 | 56.1661 | 28.5919 | 0.4850 | 53.6606 | 28.2621 | 57.1293 | 29.3581 | 0.3741 |
| **TrabSMI** | 2.7430 | 0.5283 | 2.7709 | 0.5698 | 0.4465 | 2.6679 | 0.4282 | 2.7402 | 0.5416 | 0.3564 |
| **TrabTbN** | 3.5315 | 0.4460 | 3.5921 | 0.4624 | 0.3610 | 3.4308 | 0.2863 | 3.5020 | 0.5539 | 0.3526 |
| **TrabTbTh** | 0.0461 | 0.0059 | 0.0452 | 0.0073 | 0.3740 | 0.0458 | 0.0084 | 0.0448 | 0.0063 | 0.3637 |
| **TrabTbSp** | 0.2821 | 0.0393 | 0.2790 | 0.0388 | 0.4144 | 0.2839 | 0.0349 | 0.2895 | 0.0530 | 0.3779 |
| **TrabDenTV** | 108.5440 | 29.2006 | 110.3985 | 33.2963 | 0.4382 | 109.1494 | 28.9669 | 105.4786 | 28.9701 | 0.3669 |
| **TrabDenBV** | 886.8874 | 5.7022 | 886.4616 | 30.6755 | 0.4838 | 884.9169 | 35.8608 | 881.0973 | 26.2335 | 0.3681 |
| **TrabBMC** | 0.2630 | 0.1159 | 0.2787 | 0.1619 | 0.3883 | 0.2661 | 0.1192 | 0.2593 | 0.1260 | 0.4408 |
| **TotTV** | 6.9588 | 0.8513 | 6.9316 | 0.8630 | 0.4661 | 6.8234 | 0.8568 | 6.9162 | 0.9202 | 0.3910 |
| **TotBV** | 2.7765 | 0.4416 | 2.7712 | 0.5108 | 0.4884 | 2.7066 | 0.4285 | 2.7280 | 0.4424 | 0.4480 |
| **TotBVTV** | 0.3978 | 0.0289 | 0.3982 | 0.0404 | 0.4901 | 0.3958 | 0.0307 | 0.3941 | 0.0349 | 0.4482 |
| **TotDenTV** | 403.0243 | 43.6309 | 415.1185 | 42.4080 | 0.2290 | 402.0309 | 46.9481 | 398.6980 | 52.0043 | 0.4296 |
| **TotDenBV** | 1009.3958 | 51.4607 | 1010.3157 | 58.4492 | 0.4825 | 1013.7984 | 55.2158 | 1008.8521 | 62.0656 | 0.4123 |
| **TotBMC** | 2.8099 | 0.4951 | 2.8054 | 0.5557 | 0.4909 | 2.7511 | 0.4926 | 2.7578 | 0.4974 | 0.4857 |
| **CortTV** | 6.9588 | 0.8513 | 6.9316 | 0.8630 | 0.4661 | 6.8234 | 0.8568 | 6.9162 | 0.9202 | 0.3910 |
| **CortBV** | 2.4771 | 0.3185 | 2.4614 | 0.3572 | 0.4514 | 2.4120 | 0.3132 | 2.4349 | 0.3091 | 0.4218 |
| **CortBVTV** | 0.3562 | 0.0218 | 0.3555 | 0.0330 | 0.4778 | 0.3538 | 0.0226 | 0.3535 | 0.0300 | 0.4862 |
| **CortBMC** | 2.7037 | 0.2014 | 2.5267 | 0.4255 | 0.1228 | 2.4850 | 0.3849 | 2.4985 | 0.3906 | 0.4628 |
| **CortBMD** | 1.0253 | 0.0572 | 1.0238 | 0.0616 | 0.4723 | 1.0278 | 0.0596 | 1.0234 | 0.0662 | 0.4264 |
| **MSMeanpMOI** | 0.2884 | 0.0431 | 0.4183 | 0.1371 | **0.0050** | 0.2958 | 0.0583 | 0.3915 | 0.1439 | **0.0265** |
| **MSMeanImax** | 0.1922 | 0.0335 | 0.2734 | 0.0897 | **0.0071** | 0.1969 | 0.0477 | 0.2550 | 0.0952 | **0.0394** |
| **MSMeanImin** | 0.0962 | 0.0101 | 0.1449 | 0.0480 | **0.0029** | 0.0989 | 0.0122 | 0.1365 | 0.0494 | **0.0126** |
| **MSMeanBArea** | 0.7389 | 0.1507 | 0.8403 | 0.1673 | 0.0515 | 0.7177 | 0.1366 | 0.8108 | 0.1768 | 0.0682 |
| **MSMeanTArea** | 1.5882 | 0.0745 | 1.8721 | 0.2837 | **0.0033** | 1.6067 | 0.0945 | 1.8077 | 0.2907 | **0.0212** |
| **MsTbTh** | 0.1741 | 0.0250 | 0.1878 | 0.0262 | 0.0819 | 0.1704 | 0.0256 | 0.1842 | 0.0273 | 0.0860 |
| **MsDenTV** | 480.3975 | 64.7579 | 503.2163 | 69.3336 | 0.1867 | 476.3299 | 71.8204 | 499.1016 | 63.8728 | 0.1806 |
| **MSDenBV** | 1099.4069 | 53.5213 | 1118.8063 | 55.4596 | 0.1747 | 1100.5590 | 53.1031 | 1113.4643 | 48.4555 | 0.2449 |
| **MSDenBVTV** | 2.3150 | 0.2243 | 2.2518 | 0.2235 | 0.2269 | 2.3450 | 0.2587 | 2.2568 | 0.2195 | 0.1566 |

Supplemental Table 3. Full microCT data for female *Rapsn* cKO mice tibias. P-values <0.05 are bolded.

| Female Rapsn Tibia | | | | | | | | | | |
|---|---|---|---|---|---|---|---|---|---|---|
| | Non-loaded | | | | | Loaded | | | | |
| | Cre- | St Dev | Cre+ | St Dev | p Value | Cre- | St Dev | Cre+ | St Dev | p Value |
| **TrabTV** | 1.9602 | 0.2705 | 2.1879 | 0.2255 | 0.0845 | 1.9125 | 0.3469 | 2.0858 | 0.1931 | 0.1975 |
| **TrabBV** | 0.1320 | 0.0549 | 0.1486 | 0.0471 | 0.3044 | 0.1202 | 0.0386 | 0.1472 | 0.0383 | 0.1538 |
| **TrabBVTV** | 0.0556 | 0.0136 | 0.0645 | 0.0166 | 0.1887 | 0.0598 | 0.0121 | 0.0671 | 0.0117 | 0.1866 |
| **TrabConnD** | 22.6098 | 14.2895 | 21.0933 | 6.2925 | 0.4159 | 17.5812 | 8.2846 | 17.2862 | 5.4826 | 0.4793 |
| **TrabSMI** | 2.7058 | 0.2547 | 2.5714 | 0.1894 | 0.1780 | 2.6393 | 0.4351 | 2.6077 | 0.1930 | 0.4487 |
| **TrabTbN** | 2.6103 | 0.5843 | 2.5853 | 0.4533 | 0.4698 | 2.5817 | 0.4414 | 2.6665 | 0.4487 | 0.3875 |
| **TrabTbTh** | 0.0535 | 0.0051 | 0.0530 | 0.0019 | 0.4258 | 0.0514 | 0.0055 | 0.0546 | 0.0060 | 0.2033 |
| **TrabTbSp** | 0.4006 | 0.0903 | 0.3977 | 0.0751 | 0.4783 | 0.3977 | 0.0621 | 0.3879 | 0.0684 | 0.4097 |
| **TrabDenTV** | 95.9943 | 25.1806 | 92.2053 | 16.5538 | 0.3902 | 92.3195 | 14.3621 | 95.4124 | 12.7290 | 0.3684 |
| **TrabDenBV** | 963.5983 | 22.8280 | 951.1600 | 30.0470 | 0.2273 | 952.9167 | 35.2808 | 940.7306 | 27.9370 | 0.2899 |
| **TrabBMC** | 0.1097 | 0.0376 | 0.1420 | 0.0478 | 0.1347 | 0.1146 | 0.0379 | 0.1389 | 0.0383 | 0.1757 |
| **TotTV** | 3.6907 | 0.2825 | 4.0268 | 0.2114 | **0.0281** | 3.7941 | 0.3479 | 3.9734 | 0.1541 | 0.1837 |
| **TotBV** | 1.5844 | 0.0877 | 1.6795 | 0.0946 | 0.0589 | 1.6639 | 0.0696 | 1.7221 | 0.0678 | 0.1137 |
| **TotBVTV** | 0.4304 | 0.0239 | 0.4291 | 0.0215 | 0.4652 | 0.4308 | 0.0343 | 0.4335 | 0.0110 | 0.4431 |
| **TotDenTV** | 439.7751 | 25.2086 | 420.4270 | 8.5072 | 0.0690 | 449.6032 | 39.2775 | 436.0841 | 13.9495 | 0.2669 |
| **TotDenBV** | 1044.4635 | 11.7628 | 1030.3887 | 6.5766 | **0.0209** | 1035.6995 | 10.6024 | 1028.2763 | 19.2759 | 0.2416 |
| **TotBMC** | 1.6550 | 0.0984 | 1.7308 | 0.1048 | 0.1239 | 1.7323 | 0.0789 | 1.7715 | 0.0968 | 0.2500 |
| **CortTV** | 3.6907 | 0.2825 | 4.0268 | 0.2114 | **0.0281** | 3.7941 | 0.3479 | 3.9734 | 0.1541 | 0.1837 |
| **CortBV** | 1.4524 | 0.0752 | 1.5309 | 0.0543 | **0.0418** | 1.5437 | 0.0672 | 1.5749 | 0.0376 | 0.2146 |
| **CortBVTV** | 0.3946 | 0.0224 | 0.3961 | 0.0177 | 0.4543 | 0.3805 | 0.0078 | 0.3966 | 0.0100 | **0.0149** |
| **CortBMC** | 1.5271 | 0.0774 | 1.5889 | 0.0645 | 0.0949 | 1.6177 | 0.0752 | 1.6326 | 0.0685 | 0.3795 |
| **CortBMD** | 1.0515 | 0.0129 | 1.0377 | 0.0069 | **0.0304** | 1.0479 | 0.0149 | 1.0363 | 0.0193 | 0.1574 |
| **MSMeanpMOI** | 0.2224 | 0.0309 | 0.2385 | 0.0328 | 0.2116 | 0.2220 | 0.0341 | 0.2487 | 0.0324 | 0.1257 |
| **MSMeanImax** | 0.1567 | 0.0240 | 0.1678 | 0.0281 | 0.2471 | 0.1510 | 0.0286 | 0.1694 | 0.0253 | 0.1638 |
| **MSMeanImin** | 0.0657 | 0.0080 | 0.0707 | 0.0066 | 0.1485 | 0.0711 | 0.0058 | 0.0794 | 0.0107 | 0.0738 |
| **MSMeanBArea** | 0.6906 | 0.0797 | 0.7274 | 0.0468 | 0.1943 | 0.7191 | 0.0431 | 0.7642 | 0.0314 | 0.0560 |
| **MSMeanTArea** | 1.1114 | 0.1081 | 1.1833 | 0.0560 | 0.1069 | 1.1585 | 0.0701 | 1.2306 | 0.0766 | 0.0815 |
| **MsTbTh** | 0.1938 | 0.0022 | 0.1914 | 0.0026 | 0.0761 | 0.1940 | 0.0076 | 0.2043 | 0.0064 | **0.0293** |
| **MsDenTV** | 691.1798 | 33.5107 | 698.7578 | 9.4896 | 0.3197 | 698.7018 | 25.6180 | 707.5546 | 28.2118 | 0.3102 |
| **MSDenBV** | 1135.7354 | 22.5326 | 1146.7184 | 14.4004 | 0.1865 | 1143.4108 | 19.0735 | 1143.3604 | 16.1698 | 0.4984 |
| **MSDenBVTV** | 1.6249 | 0.0623 | 1.6257 | 0.0748 | 0.4926 | 1.6346 | 0.0308 | 1.6172 | 0.0424 | 0.2360 |

Supplemental Table 4. Full microCT data for male *Rapsn* cKO mice tibias. P-values <0.05 are bolded.

| Male Rapsn Tibia | | | | | | | | | | |
|---|---|---|---|---|---|---|---|---|---|---|
| | Non-loaded | | | | | Loaded | | | | |
| | Cre- | St Dev | Cre+ | St Dev | p Value | Cre- | St Dev | Cre+ | St Dev | p Value |
| **TrabTV** | 2.4026 | 0.3204 | 2.3427 | 0.4279 | 0.3946 | 2.4743 | 0.2303 | 2.4823 | 0.4217 | 0.4831 |
| **TrabBV** | 0.3299 | 0.1142 | 0.3508 | 0.1573 | 0.3984 | 0.3626 | 0.0851 | 0.3717 | 0.1593 | 0.4489 |
| **TrabBVTV** | 0.1316 | 0.0362 | 0.1416 | 0.0437 | 0.3413 | 0.1434 | 0.0255 | 0.1425 | 0.0466 | 0.4823 |
| **TrabConnD** | 69.8298 | 26.8986 | 74.0433 | 25.0327 | 0.3997 | 77.9059 | 19.4474 | 91.3457 | 36.5511 | 0.2074 |
| **TrabSMI** | 2.1515 | 0.3227 | 2.2320 | 0.5898 | 0.3807 | 2.0913 | 0.2472 | 2.2357 | 0.5547 | 0.2678 |
| **TrabTbN** | 3.8880 | 0.7588 | 4.2258 | 0.1513 | 0.2046 | 4.1244 | 0.4636 | 4.3244 | 0.2007 | 0.2205 |
| **TrabTbTh** | 0.0544 | 0.0054 | 0.0551 | 0.0050 | 0.4137 | 0.0543 | 0.0060 | 0.0535 | 0.0046 | 0.4143 |
| **TrabTbSp** | 0.2405 | 0.0380 | 0.2292 | 0.0102 | 0.2903 | 0.2352 | 0.0289 | 0.2240 | 0.0123 | 0.2418 |
| **TrabDenTV** | 155.5128 | 35.4797 | 168.7518 | 34.1519 | 0.2758 | 166.9413 | 23.4559 | 168.3477 | 36.6658 | 0.4682 |
| **TrabDenBV** | 949.4272 | 14.7635 | 948.3280 | 14.8235 | 0.4541 | 953.0943 | 11.0073 | 936.5115 | 9.5104 | **0.0166** |
| **TrabBMC** | 0.3123 | 0.1120 | 0.3325 | 0.1487 | 0.3976 | 0.3426 | 0.0822 | 0.3485 | 0.1502 | 0.4648 |
| **TotTV** | 4.4957 | 0.4494 | 4.1385 | 0.5130 | 0.1212 | 4.4445 | 0.4186 | 4.3988 | 0.6112 | 0.4403 |
| **TotBV** | 2.0553 | 0.2343 | 1.9362 | 0.2759 | 0.2250 | 2.0151 | 0.2132 | 2.0366 | 0.2977 | 0.4438 |
| **TotBVTV** | 0.4573 | 0.0243 | 0.4454 | 0.0426 | 0.2729 | 0.4648 | 0.0156 | 0.4629 | 0.0220 | 0.4320 |
| **TotDenTV** | 457.3569 | 31.1309 | 468.2681 | 18.0756 | 0.2684 | 453.9219 | 37.0959 | 463.1439 | 20.6399 | 0.3291 |
| **TotDenBV** | 1024.3140 | 17.1192 | 1025.2500 | 10.0527 | 0.4613 | 1022.1348 | 17.2159 | 1023.1500 | 4.2681 | 0.4559 |
| **TotBMC** | 2.1054 | 0.2431 | 1.9842 | 0.2748 | 0.2261 | 2.0601 | 0.2234 | 2.0835 | 0.3020 | 0.4407 |
| **CortTV** | 4.4957 | 0.4494 | 4.1385 | 0.5130 | 0.1212 | 4.4445 | 0.4186 | 4.3988 | 0.6112 | 0.4403 |
| **CortBV** | 1.7254 | 0.2959 | 1.5854 | 0.1312 | 0.1981 | 1.6525 | 0.1582 | 1.6649 | 0.1607 | 0.4506 |
| **CortBVTV** | 0.3827 | 0.0365 | 0.3611 | 0.0282 | 0.1633 | 0.3843 | 0.0178 | 0.3800 | 0.0152 | 0.3460 |
| **CortBMC** | 1.7931 | 0.2973 | 1.6517 | 0.1444 | 0.1985 | 1.7175 | 0.1697 | 1.7349 | 0.1729 | 0.4354 |
| **CortBMD** | 1.0400 | 0.0178 | 1.0415 | 0.0097 | 0.4385 | 1.0391 | 0.0158 | 1.0418 | 0.0074 | 0.3775 |
| **MSMeanpMOI** | 0.3519 | 0.0683 | 0.3771 | 0.1094 | 0.3151 | 0.3677 | 0.0754 | 0.3504 | 0.1109 | 0.3768 |
| **MSMeanImax** | 0.2420 | 0.0511 | 0.2656 | 0.0802 | 0.2720 | 0.2606 | 0.0576 | 0.2451 | 0.0805 | 0.3534 |
| **MSMeanImin** | 0.1100 | 0.0200 | 0.1116 | 0.0295 | 0.4567 | 0.1071 | 0.0198 | 0.1053 | 0.0311 | 0.4514 |
| **MSMeanBArea** | 0.8659 | 0.0707 | 0.8807 | 0.1104 | 0.3910 | 0.8823 | 0.0848 | 0.8595 | 0.1170 | 0.3527 |
| **MSMeanTArea** | 1.4409 | 0.1427 | 1.4426 | 0.2314 | 0.4937 | 1.4652 | 0.1483 | 1.4206 | 0.2401 | 0.3478 |
| **MsTbTh** | 0.2033 | 0.0082 | 0.1983 | 0.0088 | 0.1764 | 0.1994 | 0.0061 | 0.2013 | 0.0075 | 0.3355 |
| **MsDenTV** | 677.8277 | 28.9191 | 679.8968 | 30.9834 | 0.4556 | 679.3148 | 35.8131 | 676.4052 | 25.4580 | 0.4443 |
| **MSDenBV** | 1129.3192 | 11.9244 | 1115.9750 | 3.6491 | **0.0289** | 1129.5216 | 13.3430 | 1117.1500 | 10.1635 | 0.0682 |
| **MSDenBVTV** | 1.6683 | 0.0615 | 1.6947 | 0.0361 | 0.2267 | 1.6407 | 0.0792 | 1.6532 | 0.0561 | 0.3768 |

SI Table 5. Mixed linear model results for dynamic histomorphometry data. The first column states which variables were directly compared to create the p-values for each measurement and genotype. P-value <0.05 are bolded.

| | Genotype | Ps.MS/BS | Ps.MAR | Ps.BFR/BS | Es.MS/BS | Es.MAR | Es.BFR/BS |
|---|---|---|---|---|---|---|---|
| *Male vs Female* | *Chrna1* ΔOcy/ΔOcy | 0.6881 | 0.9978 | 0.9999 | **0.0305** | 0.2643 | 0.1726 |
| | *Chrna1 +/+* | **0.0228** | 0.2 | 0.1647 | 0.4198 | 0.8998 | 0.7645 |
| | *Rapsn* ΔOcy/ΔOcy | 0.8298 | 0.5724 | 0.9999 | **<0.001** | **0.0183** | **0.0007** |
| | *Rapsn +/+* | 0.2684 | 0.7515 | 0.4549 | 0.3043 | 0.9362 | 0.5299 |
| *Cre + vs Cre -* | Female *Chrna1* | 0.3792 | 0.9211 | 0.6927 | 0.3766 | 0.8063 | 0.8144 |
| | Male *Chrna1* | 0.22 | **0.029** | 0.0943 | 0.0844 | 0.4765 | 0.291 |
| | Female *Rapsn* | 0.1349 | 0.5555 | 0.1343 | 0.3923 | 0.4372 | 0.76 |
| | Male *Rapsn* | 0.2408 | 0.367 | 0.8622 | **0.0001** | 0.1107 | **0.0159** |

SI Table 6. Mixed linear model results for mechanical testing data. The first column states which variables were directly compared to create the p-values for each measurement and genotype. P-value <0.05 are bolded

| | | Right Femur | Max force | Ultimate Energy | Failure Energy | Fracture Force | Fractur Disp | Stiffness | Yield Force | Yield Disp | PYD | Yield Energy |
|---|---|---|---|---|---|---|---|---|---|---|---|---|
| *Male vs Female* | *Chrna1* ΔOcy/ΔOcy | 0.8036 | **0.0016** | 0.4412 | 0.0049 | 0.9993 | 0.0717 | 0.3342 | 0.3128 | 0.9832 | 0.0816 | 0.913 |
| | *Chrna1 +/+* | 0.1279 | **0.0021** | 0.8311 | 0.951 | **0.0028** | 0.3577 | 0.1446 | 0.0003 | 0.972 | 0.2499 | 0.1319 |
| | *Rapsn* ΔOcy/ΔOcy | 0.1035 | **<0.0001** | **0.018** | 0.8882 | **0.0007** | 1 | **0.0001** | **<0.0001** | 0.9985 | 0.9999 | **0.0006** |
| | *Rapsn +/+* | **0.0024** | **0.0011** | 0.0765 | 0.9564 | 0.1136 | 0.9999 | 0.2262 | **<0.0001** | 0.9899 | 1 | **0.0028** |
| | WT | 0.7749 | 0.1287 | **0.0132** | **0.0023** | 0.1924 | **0.0044** | 0.8637 | 0.1083 | 0.9754 | **0.0065** | **0.0075** |
| *Cre + vs Cre -* | Female *Chrna1* | 0.6394 | 0.9337 | 0.609 | 0.7834 | 0.7214 | 0.7282 | 0.7281 | 0.3416 | 0.5821 | 0.9791 | 0.6656 |
| | Male *Chrna1* | 0.4953 | 0.8273 | 0.8107 | **0.0002** | **0.0008** | **0.0002** | 0.99 | 0.2581 | 0.5529 | **<0.0001** | 0.3906 |
| | Female *Rapsn* | 0.933 | 0.826 | 0.4599 | 0.8377 | 0.6194 | 0.9188 | 0.1944 | 0.228 | 0.5554 | 0.7222 | 0.6408 |
| | Male *Rapsn* | 0.1928 | **0.021** | 0.1682 | 0.6474 | 0.2016 | 0.7907 | 0.1584 | 0.2939 | 0.8818 | 0.7965 | 0.9043 |

SI Table 7. Mixed linear model results for MT3 osteocyte loading data. The first column states which variables were directly compared to create the p-values for each measurement and genotype. P-value <0.05 are bolded.

| | **Microstrain** | **Genotype** | **Percent Responding Cells** | **Mean Intensity** | **Max Intensity** |
|---|---|---|---|---|---|
| *Male vs Female* | 250 | *Chrna1* x GCaMP | 0.3837 | 0.7736 | 0.8179 |
| | | GCaMP | 0.9721 | 0.5467 | 0.7319 |
| | | *Rapsn* x GCaMP | 0.8002 | 0.9815 | 0.7369 |
| | 500 | *Chrna1* x GCaMP | 0.7378 | 0.6469 | 0.9156 |
| | | GCaMP | 0.273 | 0.6318 | 0.6327 |
| | | *Rapsn* x GCaMP | 0.7573 | 0.4351 | 0.1944 |
| | 1000 | *Chrna1* x GCaMP | 0.2606 | 0.8508 | 0.5401 |
| | | GCaMP | 0.0546 | 0.8054 | 0.869 |
| | | *Rapsn* x GCaMP | 0.5995 | 0.6223 | 0.631 |
| | 2000 | *Chrna1* x GCaMP | 0.737 | 0.0593 | 0.1058 |
| | | GCaMP | 0.3004 | 0.2453 | 0.2162 |
| | | *Rapsn* x GCaMP | 0.6487 | 0.089 | **0.0478** |
| | 3000 | *Chrna1* x GCaMP | 0.9117 | 0.1021 | 0.2562 |
| | | GCaMP | 0.8589 | 0.1687 | 0.2618 |
| | | *Rapsn* x GCaMP | 0.6493 | **0.007** | **0.0467** |
| *Genotype* | 250 (F) | C vs G | 0.9988 | 0.9999 | 0.9684 |
| | | C vs R | 1 | 0.974 | 0.9999 |
| | | G vs R | 0.9988 | 0.965 | 0.9564 |
| | 250 (M) | C vs G | 0.6282 | 0.9762 | 0.9797 |
| | | C vs R | 0.7003 | 0.9948 | 0.8389 |
| | | G vs R | 0.9706 | 0.9281 | 0.9189 |
| | 500 (F) | C vs G | 0.95 | 0.9621 | 0.6897 |
| | | C vs R | 0.9735 | 0.9846 | 0.8453 |
| | | G vs R | 0.9933 | 0.9911 | 0.9315 |
| | 500 (M) | C vs G | 0.484 | 0.953 | 0.8773 |
| | | C vs R | 0.9879 | 0.3092 | 0.1348 |
| | | G vs R | 0.3587 | 0.5494 | 0.4216 |
| | 1000 (F) | C vs G | 0.9581 | 0.9082 | 0.9515 |
| | | C vs R | 0.7949 | 0.9272 | 0.9685 |
| | | G vs R | 0.9241 | 0.9966 | 0.9962 |
| | 1000 (M) | C vs G | **0.0142** | 0.6621 | 0.8805 |
| | | C vs R | 0.9943 | 0.9822 | 0.6398 |
| | | G vs R | **0.0107** | 0.7213 | 0.9473 |
| | 2000 (F) | C vs G | 0.7463 | 0.8291 | 0.8473 |
| | | C vs R | 0.8098 | 0.9522 | 0.9904 |
| | | G vs R | 0.9849 | 0.9364 | 0.8788 |
| | 2000 (M) | C vs G | 0.7988 | **0.0317** | **0.0486** |
| | | C vs R | 0.7373 | 0.9534 | 0.9932 |
| | | G vs R | 0.3543 | **0.0405** | **0.0241** |
| | 3000 (F) | C vs G | 0.9284 | 0.8249 | 0.4583 |
| | | C vs R | 0.8759 | 0.7867 | 0.2984 |
| | | G vs R | 0.9934 | 0.9993 | 0.9681 |
| | 3000 (M) | C vs G | 0.9953 | **0.0363** | 0.5347 |
| | | C vs R | 0.6637 | 0.3404 | 0.0645 |
| | | G vs R | 0.7588 | **0.0003** | **0.004** |

SI Table 8. Mixed linear model results for uCT data (1 of 3). The first column states which variables were directly compared to create the p-values for each measurement and genotype. P-value <0.05 are bolded.

| | | | TrabTV | TrabBV | TrabBVTV | TrabConnD | TrabSMI | TrabTbN | TrabTbTh | TrabTbSp | TrabDenTV | TrabDenBV | TrabBMC |
|---|---|---|---|---|---|---|---|---|---|---|---|---|---|
| *Male vs Female* | Loaded | *Chrna1* ΔOcy/ΔOcy | **0.0019** | **<0.0001** | **<0.0001** | **<0.0001** | **<0.0001** | **<0.0001** | 0.0001 | **<0.0001** | **<0.0001** | **0.0108** | **<0.0001** |
| | | *Chrna1* +/+ | **<0.0001** | **<0.0001** | **<0.0001** | **<0.0001** | **<0.0001** | **<0.0001** | **<0.0001** | **<0.0001** | **<0.0001** | **0.0154** | **<0.0001** |
| | | *Rapsn* ΔOcy/ΔOcy | 0.1602 | **0.0008** | **<0.0001** | **0.0001** | 0.2596 | **<0.0001** | 0.9794 | **<0.0001** | **<0.0001** | 1 | **0.0006** |
| | | *Rapsn* +/+ | 0.7685 | 0.355 | **0.0008** | **0.0001** | 0.9274 | **<0.0001** | 1 | **0.0004** | **0.0007** | 1 | **0.0307** |
| | Non-Loaded | *Chrna1* ΔOcy/ΔOcy | **0.0001** | **<0.0001** | **<0.0001** | **<0.0001** | **0.0001** | **<0.0001** | **<0.0001** | **<0.0001** | **<0.0001** | **0.0086** | **<0.0001** |
| | | *Chrna1* +/+ | **<0.0001** | **<0.0001** | **<0.0001** | **<0.0001** | **<0.0001** | **<0.0001** | **<0.0001** | **<0.0001** | **<0.0001** | **0.0171** | **<0.0001** |
| | | *Rapsn* ΔOcy/ΔOcy | 0.4443 | **0.0132** | **<0.0001** | **0.0017** | 0.2621 | **<0.0001** | 1 | **<0.0001** | **0.0005** | 0.9794 | **0.0036** |
| | | *Rapsn* +/+ | 0.999 | 0.0796 | **0.0004** | **0.0069** | 0.9376 | **<0.0001** | 0.9992 | **0.0003** | **0.0003** | 1 | 0.0675 |
| *Cre + vs Cre-* | Loaded | Female *Chrna1* | 0.4025 | 0.9245 | 0.7788 | 0.6524 | 0.5858 | 0.2731 | 0.4734 | 0.1792 | 0.7349 | 0.7033 | 0.9317 |
| | | Male *Chrna1* | 0.6711 | 0.9692 | 0.7331 | 0.6569 | 0.9128 | 0.8209 | 0.8318 | 0.7813 | 0.687 | 0.6772 | 0.8456 |
| | | Female *Rapsn* | 0.5596 | 0.714 | 0.6683 | 0.8874 | 0.8101 | 0.8499 | 0.4503 | 0.8813 | 0.8879 | 0.2851 | 0.735 |
| | | Male *Rapsn* | 0.9795 | 0.8862 | 0.9526 | 0.3003 | 0.5856 | 0.5728 | 0.8249 | 0.7358 | 0.9258 | 0.3226 | 0.919 |
| | Non-Loaded | Female *Chrna1* | 0.7406 | 0.6982 | 0.4903 | 0.6691 | 0.937 | 0.1059 | 0.2157 | **0.0274** | 0.2795 | 0.0845 | 0.6694 |
| | | Male *Chrna1* | 0.9795 | 0.7856 | 0.8791 | 0.9576 | 0.8613 | 0.701 | 0.8632 | 0.8753 | 0.8387 | 0.5209 | 0.6562 |
| | | Female *Rapsn* | 0.3452 | 0.7918 | 0.5491 | 0.9057 | 0.6079 | 0.9239 | 0.8903 | 0.9309 | 0.7997 | 0.4421 | 0.5315 |
| | | Male *Rapsn* | 0.8058 | 0.7427 | 0.5228 | 0.7448 | 0.7612 | 0.1985 | 0.8334 | 0.825 | 0.3815 | 0.9476 | 0.7291 |
| *Load vs Non-Loaded* | ΔOcy/ΔOcy | Female *Chrna1* | 0.8723 | 1 | 1 | 1 | 0.8432 | 0.9996 | **0.0153** | 0.8889 | 0.9951 | 0.966 | 1 |
| | | Male *Chrna1* | 1 | 1 | 1 | 0.9985 | 1 | 0.9987 | 0.9941 | 1 | 1 | 0.9978 | 1 |
| | +/+ | Female *Chrna1* | 1 | 1 | 1 | 1 | 0.9993 | 0.9999 | 0.9537 | 1 | 0.9991 | 0.932 | 1 |
| | | Male *Chrna1* | 0.9918 | 0.9563 | 0.9503 | 1 | 0.9996 | 0.9246 | 0.9171 | 0.8737 | 0.9228 | 0.9042 | 0.8872 |
| | ΔOcy/ΔOcy | Female *Rapsn* | 0.9998 | 0.9999 | 0.9995 | 0.9999 | 1 | 1 | 0.7657 | 1 | 0.9997 | 0.9226 | 1 |
| | | Male *Rapsn* | 0.9931 | 0.8998 | 0.7015 | 0.8289 | 0.9989 | 0.3594 | 1 | 1 | 0.6793 | 0.9998 | 0.915 |
| | +/+ | Female *Rapsn* | 0.9678 | 1 | 1 | 0.9999 | 1 | 1 | 0.9983 | 1 | 1 | 0.7857 | 1 |
| | | Male *Rapsn* | 0.9574 | 0.999 | 1 | 0.3631 | 1 | 0.9991 | 0.9752 | 1 | 1 | 0.9539 | 0.9998 |

SI Table 9. Mixed linear model results for uCT data (2 of 3). The first column states which variables were directly compared to create the p-values for each measurement and genotype. P-value <0.05 are bolded.

| | | | **MSMeanpMOI** | **MSMeanImax** | **MSMeanImin** | **MSMeanBArea** | **MSMeanTArea** | **MSTbTh** | **MSDenTV** | **MSDenBV** | **MSDenBVTV** |
|---|---|---|---|---|---|---|---|---|---|---|---|
| *Male vs Female* | Loaded | *Chrna1* ΔOcy/ΔOcy | 1 | 1 | 0.9999 | 1 | 1 | 0.9998 | 0.8959 | 0.8046 | 0.9746 |
| | | *Chrna1* +/+ | 0.0754 | 0.1397 | **0.0078** | 0.5171 | **0.0185** | 0.897 | 0.9999 | 1 | 0.9989 |
| | | *Rapsn* ΔOcy/ΔOcy | 0.0501 | **0.0202** | 0.2919 | 0.2942 | **0.0316** | 0.9992 | 0.9995 | 0.9995 | 1 |
| | | *Rapsn* +/+ | 0.6291 | 0.5294 | 0.8531 | 0.9418 | 0.6911 | 1 | 0.9974 | 0.9888 | 1 |
| | Non-Loaded | *Chrna1* ΔOcy/ΔOcy | 0.9911 | 0.9934 | 0.9877 | 0.9999 | 1 | 0.993 | 0.8839 | 0.6527 | 0.9915 |
| | | *Chrna1* +/+ | **0.0022** | **0.0025** | **0.003** | 0.1284 | **0.0009** | 0.5963 | 0.9997 | 1 | 0.9963 |
| | | *Rapsn* ΔOcy/ΔOcy | 0.1195 | 0.15 | 0.0942 | 0.2118 | 0.0153 | 0.9898 | 0.9999 | 1 | 1 |
| | | *Rapsn* +/+ | 0.2678 | 0.2314 | 0.4089 | 0.6534 | 0.3486 | 0.9999 | 0.9999 | 0.9843 | 0.9998 |
| *Cre + vs Cre -* | Loaded | Female *Chrna1* | 0.8587 | 0.8298 | 0.7886 | 0.98 | 0.8099 | 0.9869 | 0.3886 | 0.2822 | 0.604 |
| | | Male *Chrna1* | **0.0049** | **0.0119** | **0.0009** | 0.0559 | **0.0031** | 0.1133 | 0.3714 | 0.4889 | 0.2869 |
| | | Female *Rapsn* | 0.6981 | 0.6947 | 0.7139 | 0.6492 | 0.5768 | 0.476 | 0.8251 | 0.9346 | 0.7939 |
| | | Male *Rapsn* | 0.7466 | 0.6721 | 0.9177 | 0.7761 | 0.6764 | 0.9988 | 0.945 | 0.6888 | 0.9274 |
| | Non-Loaded | Female *Chrna1* | 0.4591 | 0.3183 | 0.8597 | 0.365 | 0.705 | 0.289 | 0.2748 | 0.2364 | 0.3824 |
| | | Male *Chrna1* | **0.0004** | **0.0011** | **0.0001** | 0.0518 | **0.0001** | 0.1488 | 0.4242 | 0.3352 | 0.5016 |
| | | Female *Rapsn* | 0.7603 | 0.7575 | 0.7763 | 0.6422 | 0.4963 | 0.9488 | 0.9217 | 0.7192 | 0.9954 |
| | | Male *Rapsn* | 0.6383 | 0.5195 | 0.9283 | 0.854 | 0.9871 | 0.7281 | 0.9609 | 0.6658 | 0.8475 |
| *Load vs Non-Loaded* | ΔOcy/ΔOcy | Female *Chrna1* | 0.9583 | 0.9225 | 0.9973 | 0.8605 | 0.9535 | 0.5353 | 0.9994 | 0.9989 | 0.9999 |
| | | Male *Chrna1* | 1 | 1 | 1 | 0.9917 | 0.9994 | 0.9873 | 1 | 1 | 0.9972 |
| | +/+ | Female *Chrna1* | 0.9971 | 0.9717 | 0.9879 | 0.9998 | 1 | 0.9998 | 1 | 1 | 1 |
| | | Male *Chrna1* | 0.7245 | 0.7541 | 0.722 | 0.9194 | 0.3046 | 0.9912 | 1 | 0.9959 | 1 |
| | ΔOcy/ΔOcy | Female *Rapsn* | 1 | 1 | 0.9971 | 0.9922 | 0.9643 | 1 | 1 | 0.9955 | 1 |
| | | Male *Rapsn* | 0.9964 | 0.9377 | 0.9999 | 0.9994 | 0.9984 | 0.9999 | 1 | 1 | 0.9994 |
| | +/+ | Female *Rapsn* | 1 | 1 | 0.9954 | 0.9976 | 0.9979 | 0.5845 | 0.9996 | 1 | 0.9999 |
| | | Male *Rapsn* | 0.9893 | 0.9846 | 0.9978 | 0.9997 | 0.9999 | 0.9999 | 1 | 1 | 0.9991 |

SI Table 10. Mixed linear model results for uCT data (3 of 3). The first column states which variables were directly compared to create the p-values for each measurement and genotype. P-value <0.05 are bolded.

| | | | TotTV | TotBV | TotBVTV | TotDenTV | TotDenBV | TotBMC | CortTV | CortBV | CortBVTV | CortBMC | CortBMD |
|---|---|---|---|---|---|---|---|---|---|---|---|---|---|
| *Male vs Female* | Loaded | *Chrna1* ΔOcy/ΔOcy | **0.0001** | **0.0004** | 0.5145 | 0.8777 | 0.9997 | **0.0098** | **0.0001** | **0.0162** | 1 | 0.1217 | 0.9895 |
| | | *Chrna1* +/+ | **<0.0001** | **0.0003** | 0.9858 | 1 | 0.9979 | **0.0341** | **<0.0001** | **0.0337** | 0.65 | 0.4169 | 1 |
| | | *Rapsn* ΔOcy/ΔOcy | 0.4553 | 0.5826 | 0.7019 | 1 | 0.9998 | 0.8802 | 0.4553 | 0.996 | 0.9999 | 0.9998 | 1 |
| | | *Rapsn* +/+ | 0.9694 | 0.8784 | 0.9529 | 0.9954 | 1 | 0.9657 | 0.9694 | 0.9998 | 0.9935 | 0.9999 | 1 |
| | Non-Loaded | *Chrna1* ΔOcy/ΔOcy | **<0.0001** | **0.0001** | 0.6071 | 0.9632 | 1 | **0.0044** | **<0.0001** | **0.002** | 0.9999 | **0.0432** | 0.9996 |
| | | *Chrna1* +/+ | **<0.0001** | **0.0002** | 0.9926 | 1 | 0.9996 | **0.0226** | **<0.0001** | **0.0425** | 0.4473 | 0.4393 | 1 |
| | | *Rapsn* ΔOcy/ΔOcy | 0.1939 | 0.213 | 0.8855 | 0.9994 | 0.994 | 0.5929 | 0.1939 | 0.6083 | 0.9981 | 0.9161 | 1 |
| | | *Rapsn* +/+ | 1 | 0.9566 | 0.998 | 0.9331 | 1 | 0.9906 | 1 | 1 | 0.7887 | 1 | 1 |
| *Cre + vs Cre -* | Loaded | Female *Chrna1* | 0.8512 | 0.3581 | 0.2197 | 0.21 | 0.3404 | 0.279 | 0.8512 | 0.2734 | 0.202 | 0.2334 | 0.3423 |
| | | Male *Chrna1* | 0.6699 | 0.8693 | 0.9066 | 0.8808 | 0.8489 | 0.9685 | 0.6699 | 0.8245 | 0.9773 | 0.9267 | 0.8692 |
| | | Female *Rapsn* | 0.5857 | 0.795 | 0.8742 | 0.654 | 0.7498 | 0.9054 | 0.5857 | 0.8477 | 0.3239 | 0.9543 | 0.7627 |
| | | Male *Rapsn* | 0.8993 | 0.9205 | 0.9338 | 0.8024 | 0.9812 | 0.9329 | 0.8993 | 0.9421 | 0.8385 | 0.9428 | 0.9515 |
| | Non-Loaded | Female *Chrna1* | 0.66 | 0.2458 | 0.1993 | 0.2667 | 0.5231 | 0.2484 | 0.66 | 0.1407 | 0.1489 | 0.1851 | 0.5523 |
| | | Male *Chrna1* | 0.9005 | 0.9674 | 0.9803 | 0.8993 | 0.9717 | 0.9784 | 0.9005 | 0.8787 | 0.9608 | 0.8002 | 0.9538 |
| | | Female *Rapsn* | 0.3475 | 0.6546 | 0.9563 | 0.5957 | 0.7409 | 0.7827 | 0.3475 | 0.6415 | 0.9436 | 0.7972 | 0.7534 |
| | | Male *Rapsn* | 0.3236 | 0.5797 | 0.607 | 0.7672 | 0.9826 | 0.6629 | 0.3236 | 0.4128 | 0.3026 | 0.5614 | 0.9725 |
| *Load vs Non-Loaded* | ΔOcy/ΔOcy | Female *Chrna1* | 0.988 | 1 | 0.9925 | 0.5398 | 0.5978 | 0.9986 | 0.988 | 1 | 0.9954 | 0.9996 | 0.4151 |
| | | Male *Chrna1* | 0.5896 | 0.2255 | 1 | 1 | 0.9396 | 0.5434 | 0.5896 | 0.4787 | 0.9999 | 0.4307 | 0.9985 |
| | +/+ | Female *Chrna1* | 1 | 0.8987 | 0.982 | 0.9259 | 0.999 | 0.9583 | 1 | 0.8923 | 0.9477 | 0.9333 | 0.9995 |
| | | Male *Chrna1* | 1 | 0.3885 | 0.9816 | 0.4767 | 0.9993 | 0.3649 | 1 | 0.9402 | 0.9996 | 0.9396 | 1 |
| | ΔOcy/ΔOcy | Female *Rapsn* | 0.9962 | 0.4406 | 1 | 0.7942 | 0.9897 | 0.5808 | 0.9662 | 0.4294 | 0.4816 | 0.5202 | 0.9976 |
| | | Male *Rapsn* | 0.9988 | 0.9268 | 0.9639 | 0.9986 | 0.9997 | 0.91 | 0.9988 | 0.5387 | 1 | 0.5676 | 1 |
| | +/+ | Female *Rapsn* | 1 | 0.9894 | 0.9994 | 0.7403 | 0.9972 | 0.9975 | 1 | 0.9892 | 1 | 0.995 | 0.9998 |
| | | Male *Rapsn* | 0.3977 | 0.4003 | 0.6658 | 0.998 | 1 | 0.5199 | 0.3977 | 0.8126 | 0.5319 | 0.8242 | 1 |

Supplemental File 1: Excel file containing ANOVA test results, including post-hoc analyses.
